# Deep sea anaerobic microbial community couples the degradation of insoluble chitin to extracellular electron transfer

**DOI:** 10.1101/2025.06.30.662270

**Authors:** Yamini Jangir, Yongzhao Guo, Stephanie Connon, Sammy Pontrelli, Fabai Wu, Julia Schwartzman, Sujung Lim, Uwe Sauer, Otto X. Cordero, Victoria J. Orphan

**Affiliations:** Division of Geological and Planetary Sciences, California Institute of Technology, Pasadena, CA 91125, USA; Division of Biology and Biological Engineering, California Institute of Technology, Pasadena, CA 91125, USA; Institute of Molecular Systems Biology, ETH Zürich, Zurich 8093, Switzerland; Department of Biology, ETH Zürich, Zurich 8093, Switzerland; Department of Civil and Environmental Engineering, Massachusetts Institute of Technology, Cambridge, MA 02139, USA

## Abstract

Chitin is a major structural component of arthropod exoskeletons, and an important carbon and nitrogen source in marine environments. In anoxic sediments, its degradation generates chitooligosaccharides and N-acetylglucosamine (GlcNAc), which are fermented into smaller organic molecules and oxidized anaerobically using soluble electron acceptors or insoluble ones such as metal oxides. To date, many aspects of chitin degradation in deep-sea anoxic sediments have been overlooked, including the potential coupling of insoluble chitin degradation to metal oxide reduction, the involvement of extracellular electron transfer (EET), and the spatial organization of the microorganisms involved. Using anoxic deep-sea sediments recovered from a whale fall site, we developed an innovative workflow based on electrochemical reactors, to characterize chitin degradation in these environments. Sediment samples enriched on poorly crystalline iron oxides, and subsequently transferred into an electrochemical reactor poised at +0.22 V vs SHE, showed active anodic current production when supplied with chitin, which increased 2-fold when amended with GlcNAc. Chitin reactors were dominated by *Vallitalea* (Firmicutes), *Spirochaetota*, *Gammaproteobacteria* and *Desulfobacterota*. Exoenzyme activity assays, metabolite profiling, and continued anodic current production confirmed ongoing chitin degradation linked to EET. We observed metabolic associations between chitin degraders and secondary consumers using *in situ* imaging (16S rRNA gene FISH coupled with BONCAT and nanoSIMS). These microbial partners, within the electrode-attached community, required close proximity to the poised electrode (≤ 10 µm) to remain metabolically active. Supporting these observations, cultured isolates of *Vallitalea* sp. and *Trichloromonas* sp. recovered from the whale fall site exhibited chitin degradation and electrochemical activity, respectively. When co-cultured in an bioelectrochemical reactor, the acetate produced by *Vallitalea* sp. during chitin degradation fueled *Trichloromonas* sp., which facilitated EET, hereby demonstrating that syntrophic interactions are used to couple anoxic chitin degradation to EET in deep-sea sediments. These findings exemplify the interspecies interactions and resource optimization occurring in hard-to-reach and largely unknown deep-sea ecosystems.

## Introduction

Chitin, the most abundant natural nitrogen-containing insoluble biopolymer, is a linear chain of N-acetyl-2-amino-2-deoxy-D-glucose (GlcNAc) monomers linked by glycosidic bonds^1,2^, that forms a structural extracellular matrix in many terrestrial and marine organisms^3–5^. It is synthesized in the ocean at a remarkable rate of approximately at 10^12^–10^14^ tons annually and is the second most abundant component of marine snow, particulate matter that is largely recalcitrant to microbial remineralization^2,3^. Despite its continuous deposition onto marine sediments, very little (<1%) accumulates in oxic deep-sea sediments, due to efficient microbial degradation and turnover^4–7^. Microbial chitin degradation is a vital part of the biogeochemical cycling of carbon and nitrogen and it is facilitated by extracellular enzymes that detect, bind, and cleave chitin into smaller soluble oligomers or monomers, via the hydrolytic^6^ or oxidative^8,9^ pathways. Both of these produce diverse metabolic byproducts that promote community dynamics such as cross-feeding, and influence microbial ecology and community dynamics. Generally speaking, chitin degradation and subsequent mineralization involves a complex interplay between primary chitin degraders and secondary consumers (non-degrading community members).

Importantly, the absence of a metabolically active secondary consumer can slow down the degradation process due to the accumulation of inhibitory byproducts^10,11^, such as N-acetylglucosamine, glucose, and acetate. This inhibition can be alleviated by the presence of an inexhaustible terminal electron acceptor that facilitates constant removal of inhibitory byproducts through respiration by the secondary consumer.

Chitin degradation has been extensively studied in the oxic coastal water column^1,5,6^, with increasing understanding of the responsible microbial communities^12^, as well as their functional roles^13–15^ and spatial organization^16^. Conversely, the microbial communities and subsequent processes governing this breakdown is largely unknown in anoxic deep-sea sediments, despite the prevalence of chitin degradation in these environments^17–20^. Studying chitin degradation at the microbial community level is particularly difficult in deep-sea sediment, because of the highly heterogeneous nature of these substrates and the challenges inherent to sampling and studying deep-sea microorganisms under laboratory conditions. Specifically, reproducing natural variations in the distribution of chitin particles, microbial communities, pH, temperature, and redox potential is a highly complex endeavor.

Theoretically, under the anoxic conditions that characterize deep-sea sediments, chitin degradation should be driven by terminal electron acceptors such as sulfate and CO_2_, as well as insoluble redox-active minerals containing sulfur, manganese, and iron^21^. Microbes that utilize insoluble minerals such as iron or manganese oxides as electron acceptors employ extracellular electron transfer (EET) mechanisms^22,23^. These microorganisms can be enriched and studied using electrochemical methods with soluble electron donors, where poised electrodes set at appropriate reduction potentials mimic natural electron sinks^24,25^ to act as inexhaustible terminal electron acceptor. Electrochemical approaches have proven invaluable for cultivating and characterizing electroactive microbial communities, both *in situ* and *ex situ*, expanding our understanding of their diversity and metabolic capabilities^26,27^. Indeed, measuring substrate dynamics electrochemically, in real-time, alongside microbial EET, provides a direct link between metabolic activity and current generation. Yet, while previous studies have explored electrochemical degradation of particulate matter, including chitin, in wastewater systems^28,29^, the potential for such mechanisms to facilitate chitin degradation in anoxic deep-sea sediments remains unknown.

To characterize the black box surrounding insoluble chitin polymer degradation in these environments, we examined whether syntrophic interactions occurred between chitin degraders and EET-capable organisms in anoxic chitin-rich marine sediments. We further posited that microbial communities capable of coupling the respiration of insoluble carbon source to anodic growth would display specific spatial organizations reflecting a division of labor. To test these hypotheses, we implemented an innovative workflow, adapting the use of electrochemical systems to enrich and monitor chitin degradation dynamics and to investigate the ecophysiology of responsible microbial community members recovered from deep-sea sediments.

Our approach involved collecting deep-sea sediments from a whale fall site (WF1018) within the Monterey Canyon, in the oxygen minimum zone at a depth of 1,018 meters^30^. The site hosts macrofauna such as polychaetes and crustaceans, as well as microbiomes of mesopelagic organisms^30^, making it a natural reservoir of chitin in anoxic deep-sea environments^31–33^. Using long-term electrochemical incubations under anoxic conditions, we enriched microbial communities capable of both chitin degradation and electrode respiration. Our system revealed the importance of both microbe-microbe and microbe-electrode interactions in shaping a stable community and demonstrated functional partitioning between chitin degradation and EET processes.

Moreover, we showed that electrochemical systems can be used not only to enrich electrode-respiring communities but also to dynamically monitor metabolite exchange (e.g., acetate) during particulate organic matter degradation under anoxic conditions. Finally, we isolated two dominant community members, *Vallitalea* sp. sp2 and *Trichloromonas* sp. sp17, and reconstituted a functional two-member community capable of coupling chitin degradation to electrode reduction, thus providing a controlled demonstration of insoluble polymer-EET integration in an anoxic marine sediment model.

## Results

### Electrochemical signature of a deep-sea sediment microbial community performing anoxic chitin degradation

The premise of this work relied on the successful enrichment of a native microbial community from deep-sea sediments capable of anaerobically degrading insoluble chitin and reducing solid-phase iron/electrodes. Chitin is an insoluble, semi-crystalline organic polymer. This structure coupled with the inherently slow kinetics of anaerobic metabolisms and spatial separation of the primary carbon source insoluble electron acceptor considerably increases incubation times for enrichment cultivation. Here, we implemented a combination of parallel and sequential enrichment strategies using deep-sea whale fall sediments as the inoculum (Figure. 1a) for long term enrichments of anaerobic chitin-degrading microorganisms coupled to iron oxide/poised electrode reduction. We specifically selected sediments from a whale fall site identified as locally enriched in chitin ^30^, within the oxygen minimum zone of the Monterey submarine canyon^30,34^. A spike in porewater iron (0.13 mM Fe^2+^) 1-2 cm below the sediment-water interface sediment-water interface (Figure 1 b), for the background core (2 m away), suggested an ample source of iron oxide in the region. The iron-rich sediment horizon collected (1-2 cm) was used as the inoculum for our anaerobic enrichments, and geochemical and microbial 16S rRNA analyses were conducted over time.

**Figure 1.**
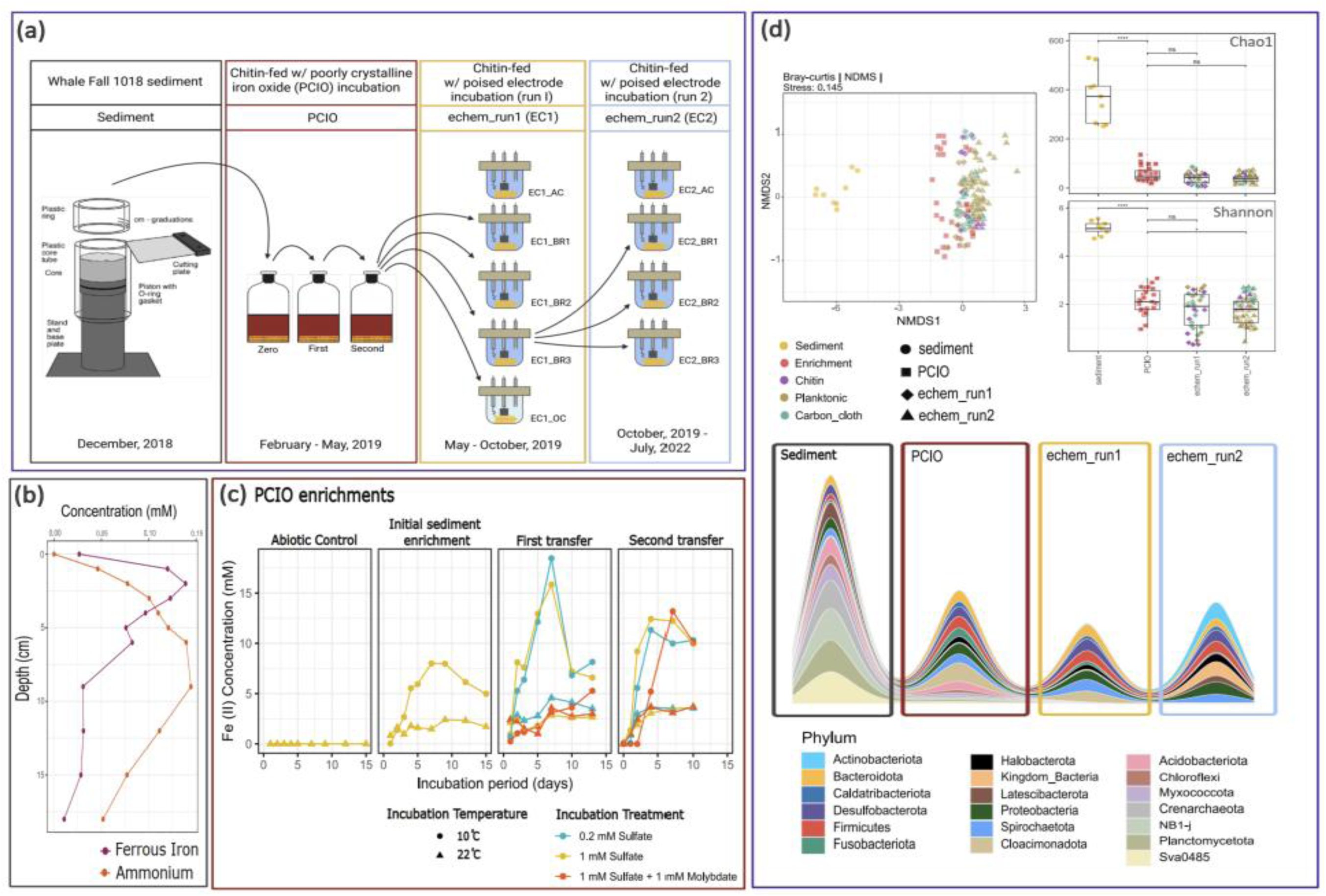
(a) Experimental workflow and microbial community analysis (a) Sediment samples (1–2 cm depth) were collected from the whalefall site WF1018 in Monterey Canyon, CA, in December 2018. These sediments were incubated with chitin (0.1 g/L) and poorly crystalline iron oxide (PCIO; 0.13 g/L). After two sequential transfers, a 5 mL aliquot of the chitin-iron enrichment was used to inoculate triplicate electrochemical reactors (BR1, BR2, and BR3) poised at +0.22 V vs SHE, along with an open circuit control (OC). This experimental setup is referred to as echem_run1 or EC1. For the second electrochemical experiment (echem_run2 or EC2), inoculum was derived from the poised electrode (carbon cloth) from EC1_BR3. (b) Sediment porewater Fe2+ and ammonium concentrations with depth collected from the whale fall site. Sediment layer (1–2 cm) chosen as the inoculum based on predicted peak in iron reduction (Fe^2+^) (c) Microbial communities enriched on chitin and PCIO were incubated at either 10°C or 22°C and with three sulfate conditions (0.2 mM SO_4_^2-^, 1 mM SO_4_^2^, and 1 mM SO ^2^ with 1 mM molybdate to inhibit sulfate reduction) and assessed for Fe^3+^ reduction via the ferrozine assay. The figure presents the results of the ferrozine assay conducted on successive PCIO enrichments. The abiotic control contains no sediment inoculum, while “zero” represents the initial sediment inoculum. “First” refers to the first transfer from the 10th-day enrichment sediment-based “zero” enrichment, and “second” indicates the transfer from the 10th-day enrichment of the “first” enrichment. Higher activity was observed at 10°C, with minimal differences across sulfate conditions. (d) (top left) NMDS ordination of Bray–Curtis community dissimilarities based on ASVs from 16S rRNA sequencing shows divergence from the native whalefall sediment community and similarity among chitin-fed iron and electrochemical incubation communities. (top-right) Alpha diversity indices reveal a decrease in species richness and evenness from native sediments to chitin-fed iron and electrochemical incubations. Stream plots show that only select microbial lineages were maintained chitin-fed iron and electrochemical incubation. here, Sediment represents whalefall samples 0-1 cm, 1-2 cm, and 2-3 cm combined, Enrichment: Chitin-PCIO enrichment (PCIO_run1; 10°C and RT, all sulfate amendments included), Chitin: Chitin-attached biomass in electrochemical incubations, Planktonic: free-living biomass in the electrochemical incubations, Carbon_cloth: Electrode-attached biomass in the electrochemical incubations]

Anaerobic microbial communities enriched with chitin and poorly crystalline iron oxide (PCIO) reduced iron, with ferrozine assays showing higher iron reduction activity at 10 °C (i.e. closer to the *in situ* temperature of 4.2 °C, compared to ambient incubation temperature 22 °C) (Figure 1c). Comparable activity levels were observed across all three sulfate conditions (0.2 mM, 1 mM, and 1 mM sulfate + 1 mM molybdate), supplied as an anabolic sulfur source (see Supplementary Materials) (Figure 1c). We next assessed the ability of these chitin-degrading, metal-reducing microbial enrichments to respire an electrode in controlled bioelectrochemical reactors, where EET respiration can be directly quantified through current production. To do so, the enriched community was used as the inoculum for the first electrochemical run (echem_run1; EC1) in three biological replicates (BR1, BR2, BR3) and one open circuit control (OC). An abiotic control (AC) was run in parallel with no inoculum. AC and BR1-3 operated with working electrodes poised at +0.22 V vs Standard Hydrogen Electrode (SHE) to mimic iron oxide reduction potential and therefore provided an electrochemical platform to allow real-time monitoring of metabolic activity after chitin amendment. In effect, EC1 served as the initial enrichment phase to select for, and stabilize, a community capable of both chitin degradation and electrode respiration under controlled anoxic conditions. Using the most electrochemically active reactor (EC1_BR3), we established the second round of electrochemical reactors (echem_run2; EC2), in series, to systematically investigate and refine the microbial community coupling chitin degradation to EET, and to assess the metabolic dynamics, substrate turnover (chitin versus monomers) and syntrophic interactions (Figure 1a). This two-step approach ensured not only the enrichment of a specialized electrogenic microbial community, but also enabled the fine-scale examination of metabolic processes, community stability, and spatial organization among interacting community members, providing robust evidence for the coupling of polymer degradation to EET in deep-sea sediment systems.

The composition of the enriched microbial community was clearly different between the *in situ* sediments and our laboratory enrichments (Non multidimensional scaling, NMDS, based on Bray-Curtis dissimilarity; stress = 0.145; Figure 1d). The enriched communities accounted for 23.1% of the observed variation (PERMANOVA analysis with 999 permutations, R² = 0.231, p < 0.001), indicating that the enrichments retained a subset of the native community, likely involved in chitin degradation. Our poised electrochemical incubations also appeared to faithfully replicate the metal-oxide metabolisms of interest, as neither Chao1 nor Shannon indices, or NMDS analyses showed significant differences between the iron and electrochemical enrichments (Figure 1d, top right). The few microbial lineages that were significantly enriched under laboratory incubations (PCIO and electrochemical) belonged to Spirochaetota, Proteobacteria, Halobacterota, Firmicutes, Desulfobacterota, and Bacteroidota (SIMPER and Indicator Species Analysis, p = 0.001, Figure 1d, bottom, and Supplementary Materials).

Having successfully enriched a microbial community capable of coupling anaerobic chitin degradation to iron oxide reduction, we tested for and confirmed the presence of active chitin degradation under anoxic conditions within EC1 (Figure. 2a). While abiotic controls (EC1_AC) exhibited no anodic current, both EC1_BR1 and EC1_BR3 did. Among the 3 replicates, EC1_BR3 showed a continuous increase in the anodic activity (current production) from 0 to 0.2 mA over a period of 2 months. Cyclic voltammetry under turnover conditions (observed in the presence of electron donor) revealed catalytic oxidation of a carbon source (sigmoidal shape) at an onset potential of-0.25 V vs SHE in both EC1_BR1 and EC1_ BR3. An additional reduction peak (0.15 V vs SHE) was observed exclusively in EC1_BR1 (Figure. 2b). None of the redox peaks were observed in the controls (EC1_AC and EC1_OC). EC1_BR3 consistently had the highest number of annotated exometabolites, with 85 identified across the incubation period, compared to 36 in EC1_BR1 and 11 in EC1_BR2 (see Supplementary Materials; Figure 2c), including a spike in GlcNAc exclusively in EC1_BR3 at day 79. Finally, acetate was fully consumed within 120 days of chitin incubation, while 5 mM residual ammonia, assumed to be sourced from chitin and GlcNAc degradation, remained (Figure 2d). Collectively, these observations indicated the presence of an anaerobic chitin-degrading microbial community capable of EET to poised electrodes and demonstrated the strength of our experimental approach.

**Figure 2.**
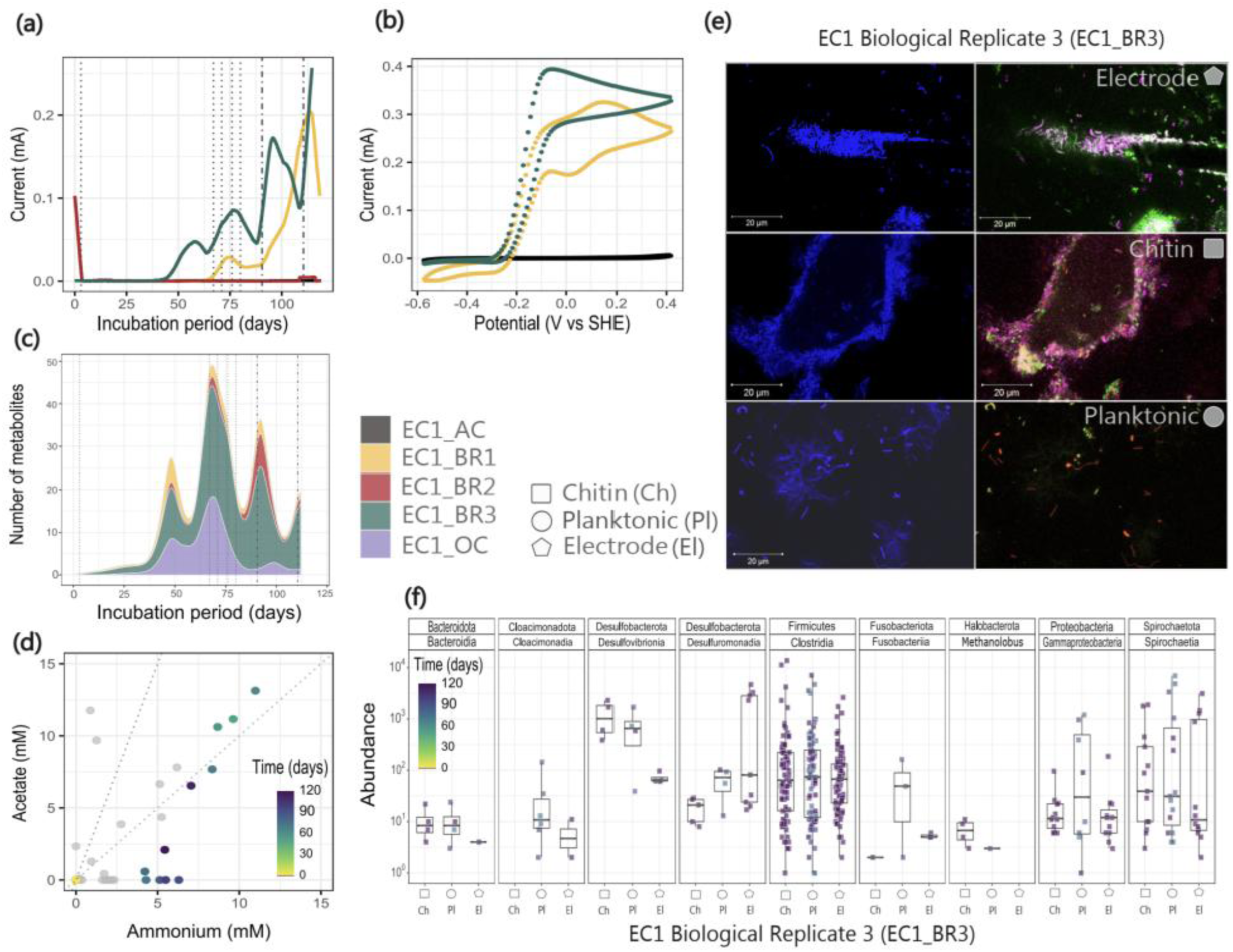
Overview of the first electrochemical reactor experiment (EC1), conducted over 120 days from May to October 2019. The figure presents data from the following setups: abiotic control (EC1_AC) in black, three biological replicates (EC1_ BR1, EC1_BR2, EC1_BR3 in yellow, red, and green respectively) and the open circuit control (EC1_OC) in purple. (a) Biological replicate 3 (EC1_BR3, green) showed the highest anodic activity. The vertical line indicates time points where samples were collected to measure the exometabolites for metabolomics and IC. Anodic current measured in the abiotic control was below 10^-5^ mA. (b) A cyclic voltammogram at a low scan rate (1 mV/s) displayed a sigmoidal curve, suggesting catalytic turnover of a carbon source. (c) Exometabolities measured by mass spectrometry for all replicates and controls are shown using stacked stream plots. BR3 produced the highest number of (putative) annotated extracellular metabolites throughout the 120-day electrochemical enrichment run. (d) Acetate and ammonia production in each biological replicate (with BR3 highlighted) is shown. Slope lines indicate acetate/ammonium ratios of 1:1 and 3:1. For EC1_BR3, FISH and 16S rRNA-based microbial community analysis was conducted for the phases—chitin-associated (Ch; square), planktonic (Pl; circle), and electrode-attached (El; pentagon). e) DAPI stained (left; blue) and 16S rRNA gene FISH images (right) of electrode-attached, chitin-attached, and planktonic biomass targeted *Gammaproteobacteria* (red), *Deltaproteobacteria* (green), and *Alphaproteobacteria* (magenta). (f) Amplicon sequence variants (ASVs) representing diverse microbial phyla and classes were identified in planktonic, chitin-attached, and electrode-attached communities. Planktonic samples were collected at four distinct time points: day 78, day 87, day 99, and day 119 of chitin incubation. The chitin-attached and electrode-attached communities were sampled on day 119.

We further demonstrated spatial partitioning of microbial taxa within the bioreactor. Indeed, different taxa were found associated with the electrode, on chitin particles, and within the planktonic phase, based on fluorescent in situ hybridization (FISH) (Figure 2e) and differential analysis of 16S rRNA amplicon sequencing data (Figure 2f and Supplementary Figure 2) Specifically, FISH probes targeting *Proteobacteria* confirmed abundance of microbial cells in EC1_BR3 after 120 days of operation, with *Gammaproteobacteria*, *Alphaproteobacteria*, and *Deltaproteobacteria* in all the three phases. Additionally, chitin particles were dominated by *Halodesulfovibrio, Roseimarinus, Methanolobus, Fusibacter, Vallitalea, Sedimispirochaeta, and Spirochateta_2,* and the electrode was enriched in *Trichloromonas (*a recently proposed clade in *Desulfuromonadaceae*^35^), and the planktonic phase included bacterial representation from *Psychromonas, Clostridia JTB215, Sphaerochaeta, and Shewanella*.

### EET supports anaerobic chitin degradation through acetate oxidation

The second set of electrochemical experiments (EC2, BR1-3) allowed for a detailed examination of microbial metabolic dynamics and substrate utilization under controlled conditions. This run was conducted with chitin as the carbon source, with replenishment of chitin during the incubation indicated; compared against separate incubations amended with soluble monomers (GlcNAc, glucose, acetate). The average anodic current, used as a proxy for metabolic activity, correlated strongly with the Shannon diversity index (Figure 3a). The microbial diversity moderately declined after removal of planktonic cells and the addition of fresh chitin, followed by stabilization of community diversity with renewed substrate availability. Anodic current production during the chitin incubations were, on average, significantly lower (∼2-fold) than that observed with the GlcNAc monomer as the carbon source (Figure 3b).

**Figure 3.**
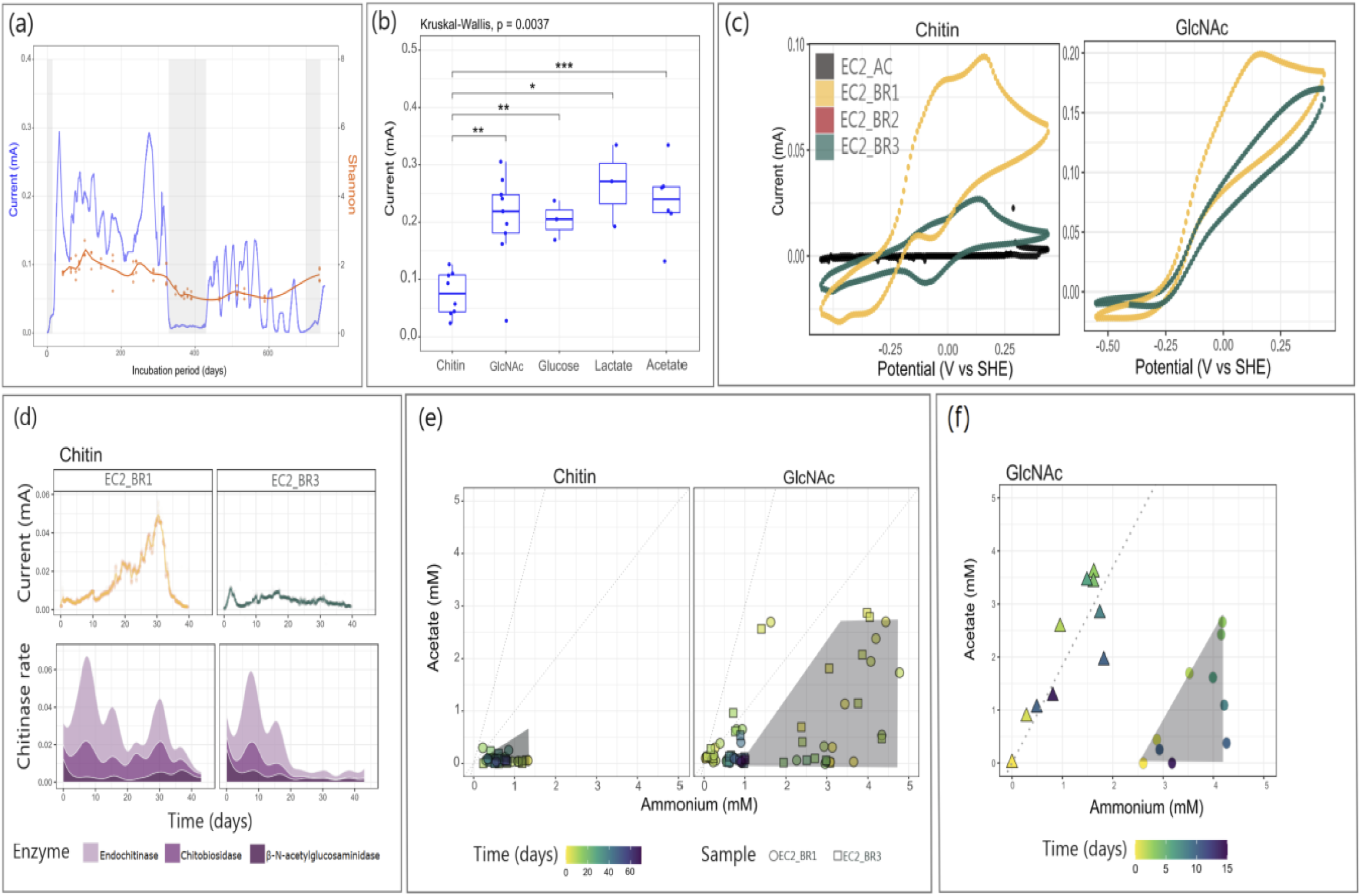
Summary of the second electrochemical reactor (echem_run2, EC2) long-term 32 months incubation (Nov 2019 – July 2022) with insoluble chitin, soluble monomer N-acetylglucosamine (GlcNAc), and other soluble organic carbon sources (lactate, pyruvate, fumarate, acetate). (a) Anodic current (metabolic activity) was observed in all three replicates (EC2_BR1-3), with a 1.4-day rolling average shown. The grey shaded regions correspond to planktonic phase removal and addition of fresh chitin. The Shannon diversity index indicates an establishment of a stable microbial diversity, even after planktonic cell removal and fresh chitin addition. Anodic current measured in the abiotic control was below 10^-5^ mA. (b) Anodic current production during chitin incubation was in comparison with GlcNAc and the carbon source. The box plot shows the average of maximum anodic current measured across the biological replicates (EC2_BR1-3) for each carbon substrate. (c) Cyclic voltammogram during chitin incubation, for all the biological replicates, showed a sigmoidal curve with a catalytic turnover potential at-0.25 V vs. SHE, a peak at 0.175 V vs. SHE, and a faint peak at-0.1 V vs. SHE (more evident for EC2_BR1). In contrast, the GlcNAc incubation showed a boost in catalytic activity with higher anodic current without additional signature peaks. (d) Temporal variation, in correspondence with the anodic current (upper) and for exochitinase activity rate (bottom; sampling frequency: 1 day) profiles, is illustrated for EC2_BR1 and EC2_BR3. Higher endochitinase activity and correlated N-acetylglucosamidase activity aligned with anodic current levels. (e) Temporal profiles of acetate and ammonia, in EC2_BR1 and EC2_BR3, revealed both lower accumulation and faster consumption during chitin incubation (0.2 g/L) compared to GlcNAc (3 mM). This pattern, as depicted in grey shaded regions, likely reflects the slower rate of anaerobic microbial chitin degradation compared to acetate and ammonia consumption, microbial growth and respiration, with time. (f) Temporal profile of average acetate versus average ammonia concentration, measured across EC2_BR1-3, during a 15 day long GlcAc incubation (circles) with corresponding incoming acetate and ammonia evaluated theoretically (triangles) using measured real-time anodic current (with coulombic efficiency, CE: 0.75), acetate assimilation rate (3.2 μM/hr), and ammonia assimilation rate (0.7 μM/hr) provides acetate:ammonia ratio as 1.84.

However, anodic responses to simpler soluble organics such as glucose, lactate, and acetate were comparable to those of GlcNAc. As observed in EC1_BR3, a steady-state catalytic turnover (with 3 mM GlcNAc) followed a classic sigmoidal curve with an onset potential of-0.25 V vs SHE, inflecting at a half-saturation potential slightly lower than −0.125 V. Notably, this onset potential is close to that of terminal multiheme cytochrome responsible for EET in *Geobacter sulfurreducens* (-0.2 V vs SHE)^36^, *Desulfuromonas soudanensis* (-0.19 V vs SHE)^37^, and *Shewanella oneidensis* (-0.23 V vs SHE)^38^ suggesting that the electrogenic microbes, presumably *Trichloromonas*, may also be using cytochromes with similar midpoint potential. During anaerobic chitin degradation, the voltammogram appeared asymmetrical and non-sigmoidal, exhibiting two prominent redox peaks with midpoint potentials of-0.125 V and +0.185 V vs SHE (Figure 3c). These observations illustrate diffusion-limited kinetics at lower substrate levels and the influence of soluble chitin degradation metabolites on EET^39,40^.

During the third phase of chitin incubation (duration ca. 50 days), EC2_BR1 and EC2_BR3, were evaluated for exochitinase activity and exometabolite profiles in relation to anodic current production. Predicted chitin metabolites, ammonia and acetate, were also monitored (Figure 3e). Similar patterns were observed between replicate bioreactors, despite differences due to variations in measured anodic current. Notably, the activity of exoenzymes, endochitinase and chitobiosidase, increased within 7 days of chitin addition, while N-acetylglucosaminidase activity remained relatively low throughout the 50-day incubation period (Figure 3d).

Exometabolite analysis via untargeted mass spectroscopy revealed two unannotated ions (m/z: 237.255, 103) that increased over time in the reactors (Supplementary Figure 3). The identity of these metabolites remains unknown at this time. Other metabolites, such as acetate, showed rapid uptake by the electrode-attached *Trichloromonas*-dominated community, indicating that anodic current was limited by the supply of the metabolites presumably produced during chitin degradation. Notably, the acetate-to-ammonia concentration ratio consistently remained below 1, suggesting rapid consumption of acetate, as each GlcNAc (a monomer of chitin) molecule is predicted to be deacylated and deaminated to produce acetate and ammonia with a ratio of 3:1 based on stoichiometry (see Supplementary Materials). While other metabolites were produced, acetate appeared to predominantly drive the activity of the electrode-attached microbial community. This was supported using a simplified mathematical model of GlcNAc metabolism to the production of acetate, followed by oxidation of acetate by electroactive microorganism on a poised electrode (See Supplementary Materials and Supplementary Figure 4). The temporal profile of measured acetate concentrations (average of EC2_BR1-3), suggests that the ratio of acetate: ammonia produced during anaerobic chitin degradation could lie anywhere between 1 to 3, most likely due to the heterogeneity within the microbial population.

To better capture dynamic metabolic behavior, we modeled the acetate-to-ammonia production ratio based on real-time current measurements rather than average current values (see Supplementary Materials) and infer that the net influx of acetate and ammonia at any time point maintained a stoichiometric ratio of approximately 1.84:1 (Figure 3F). These results support a pivotal role of EET for acetate removal in sustaining efficient anoxic chitin degradation coupled to electrode, and metal reduction, presumably. Importantly, they demonstrate that through the tracking of real-time anodic current production, these reactors can also provide an assessment of acetate flux through the community during chitin degradation.

### Spatial and temporal heterogeneity in microbial interactions

Consistent with the patterns found with EC1 reactors, the EC2_BR1-BR3 reactors also showed a clear partitioning of microbial assemblages associated with planktonic, electrode-attached, and chitin-attached phases (Figure 4). Notably, *Trichloromonas* and *Desulfuromonas* were again predominantly found in electrode-attached biomass, *Pseudomonas (Pseudomonadaceae)* were mostly found in the planktonic and chitin-attached biomass, while *Vallitalea (Firmicutes)* was evenly distributed across all phases, along with several lower-abundance taxa (e.g., *Methanolobus* and *Bacteroidetes*, Figure 4a). The strong spatial partitioning within the reactor suggested that *Pseudomonadaceae* and *Vallitalea* (planktonic and chitin-associated) are involved in primary chitin degradation and secondary metabolism, while members of the *Desulfobacterota* phylum (electrode-associated) are likely responsible for facilitating EET and presumably iron oxide reduction *in situ*.

**Figure 4.**
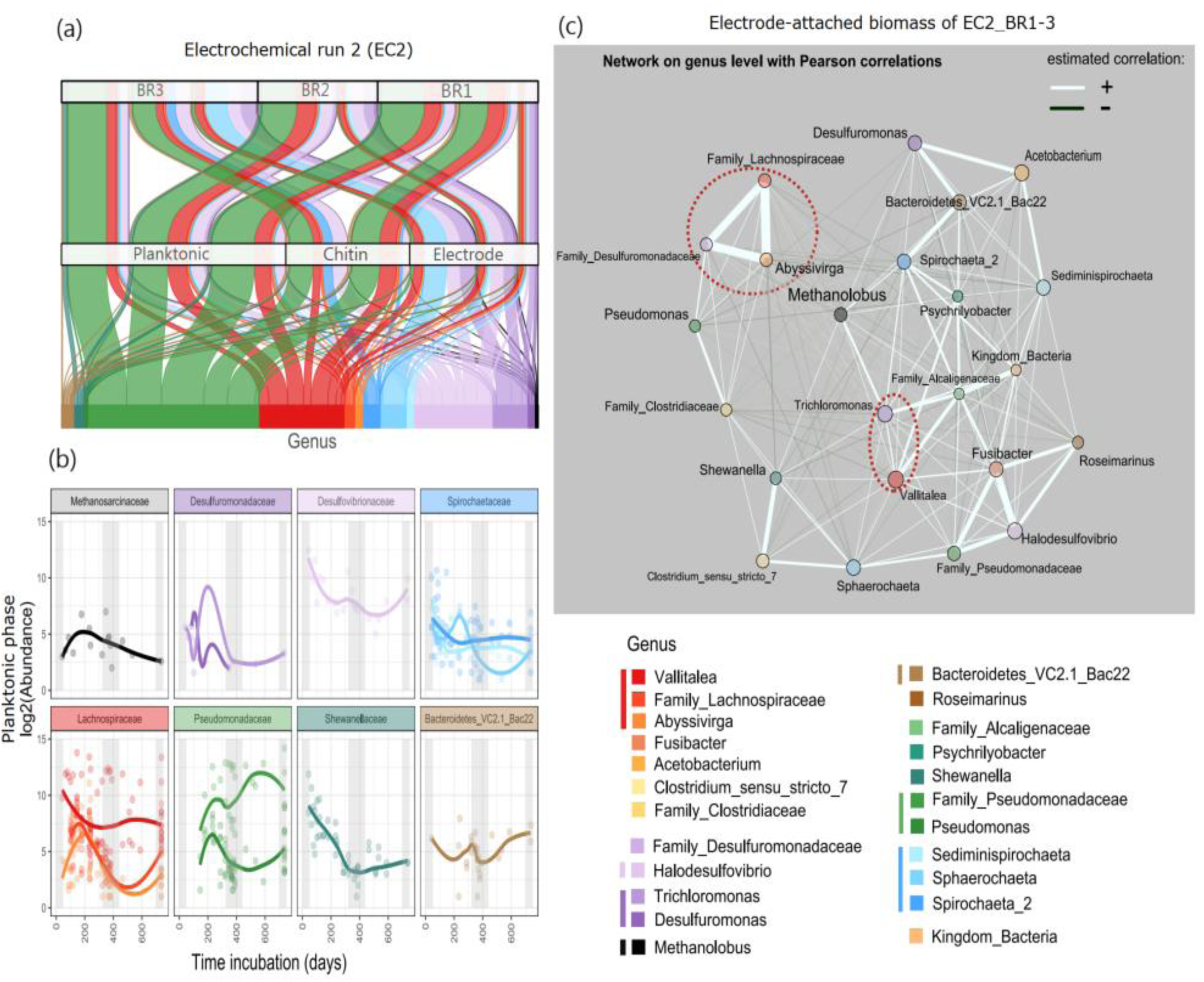
Summary of 16S rRNA gene analysis from second electrochemical run (echem_run2; EC2). (a) Amplicon sequence variants (ASVs) detected across all three biological replicates (EC2_BR1, EC2_BR2, EC2_BR3), showing relative abundances in planktonic, chitin-attached, and electrode-attached biomass. (b) Temporal dynamics of the planktonic microbial community, highlighting genera like *Sphaerochaeta, Vallitalea, Pseudomonas*, and *Shewanella* that were lost and later reestablished after refreshing with chitin on day 420 (shaded grey). (c) Compositionally aware network analysis of electrode-attached biomass in the three biological replicates, showing associations between mineral reducers (*Desulfuromonadaceae* family) and chitin degraders (*Lachnospiraceae* family). Line thickness represents the strength of the correlation. ASVs that could not be classified at the genus level but belonged to a known family were labeled using the family name.

*Vallitalea* and novel *Pseudomonadaceae* lineages represented key taxa in the planktonic phase over time and amid changes in experimental conditions in the reactor (e.g., addition of chitin and different soluble carbon monomers). *Desulfuromonas* was completely lost in the planktonic phase after 320 days of incubation and was only detected in physical association with poised electrodes thereafter (Figure 4b). Importantly, a *Vallitalea* sp. isolated from deep-sea sediments (Guaymas basin) was previously described as a potential chitin degrader^41^, and genera within the *Desulfuromonadaceae* family, capable of sulfate and metal reduction, can interact with poised electrodes by oxidizing soluble organic compounds, including acetate^42^.

A co-occurrence network analysis of the electrode-attached community (23 taxa across 11 samples; EC2_BR1-BR3) highlighted potential interactions between taxa during chitin degradation and extracellular electron transfer (Figure 4c). For example, *Vallitalea*, a dominant chitin degrader, co-occurred with *Trichloromonas*, an electroactive member closely related to *Desulfuromonas*, suggesting a potential syntrophic interaction in which fermentative byproducts like acetate fuel EET. This predicted association may further be bolstered by affiliations with other fermenters (e.g., *Lachnospiraceae*, *Acetobacterium, Pseudomonas*) and EET-capable genera (*Shewanella*). An unclassified member of *Desulfuromonadaceae*, typically involved in metal reduction, was positively correlated with both methylotrophic methanogens (*Methanolobus*) and putative fermenters, suggestive of metabolic cross-feeding. Overall, the network predicts the occurrence of multiple subclusters each comprised of putative chitin degraders, fermenters, and metal/electrode reducers, suggesting functional redundancy and raising the question of the degree of complexity required within the microbial community to fully couple insoluble chitin degradation with electrogenic activity in anoxic environments. Integrating information about the taxonomic composition across the three phases, the co-occurrence network analysis, and current predictions of metabolic potential for the dominant microorganisms, allowed us to develop the hypothesis of a metabolic link between the firmicute *Vallitalea* and members within the *Desulfuromonadaceae*.

To test this possibility and confirm the role of *Vallitalea* in the electrochemical bioreactors, we applied complementary methods using fluorescence microscopy and single-cell activity assays. Specifically, we designed a 16S rRNA FISH probe targeting *Vallitalea sp.* to visualize it in the planktonic and the electrode-attached community, and combined this taxonomy-based identification with either bioorthogonal non-canonical amino acid tagging (BONCAT), to identify translationally active microbes at the single-cell level^43^ or nanoscale secondary ion mass spectroscopy (nanoSIMS), to track and quantify the assimilation of nitrogen from ^15^N-labelled GlcNAc into cells^44^. This allowed us to implement FISH-BONCAT and FISH-nanoSIMS, previously used to link taxonomic identity to function (activity) to unravel metabolic interactions at the single-cell level^43,45–47^.

In the planktonic phase of EC2_BR3, filamentous *Vallitalea* cells were translationally active within 5 days of chitin amendment (FISH-BONCAT analysis, Figure 5a). In addition, stable isotope incubations, with 3 mM ^15^N-labeled GlcNAc, demonstrated significant ^15^N incorporation into *Vallitalea* cells by day 3, both in planktonic and electrode-attached communities indicating their ability to assimilate soluble monomers of chitin (FISH-nanoSIMS; Figure 5b). To further visualize the spatial and temporal dynamics of electrochemical and metabolic activities, we analyzed FISH and nanoSIMS images for the planktonic and the electrode-attached communities with corresponding anodic current profile (Figure 5b). Distinct phases of electrochemical (metabolic) activity were observed in the bioreactor over time with an increase in anodic current (catabolic activity) observed within 24 hours of ^15^N-labeled GlcNAc amendment. The current levels decreased on days 3 and 12 from destructive sampling of the electrode-attached community, and subsequently stabilized at approximately 0.06 mA (on day 3) and 0.05 mA (day 12). Between days 15 and 20, the current gradually declined to nearly 0 mA, indicating electron donor depletion. In the planktonic phase, during the early time point (ca. day 1.5), ^15^N assimilation from GlcNAc was minimal in all cells, consistent with low current and limited metabolic activity. By day 3, the ^15^N uptake detected in *Vallitalea* cells (planktonic phase and electrode-attached communities) reflected *Vallitalea’*s central role as a primary degrader and fermenter of GlcNAc, likely catalyzing the anodic response of the electrogens by increasing local acetate production. By day 12, substantial ^15^N incorporation was still measured in planktonic *Vallitalea* cells, but limited in other microbial taxa (identity unknown). Conversely, on the electrode-attached biofilm, the microbial cells surrounding *Vallitalea* exhibited elevated ^15^N enrichment, providing evidence of cross-feeding interactions. These could be fueled by nitrogen-containing byproducts of *Vallitalea’s* fermentation, beyond acetate (including NH_4_^+^), supporting growth of adjacent EET-capable microbes. These spatially resolved cell-specific activity measurements revealed the importance of local substrate availability (GlcNAc byproducts) in driving metabolic activity and current production within the electrode-attached microbial community. Overall, these results support the fact that spatial proximity between the primary degraders and EET-capable microbes to the poised electrode facilitates efficient cross feeding within the electrogenic biofilm.

**Figure 5.**
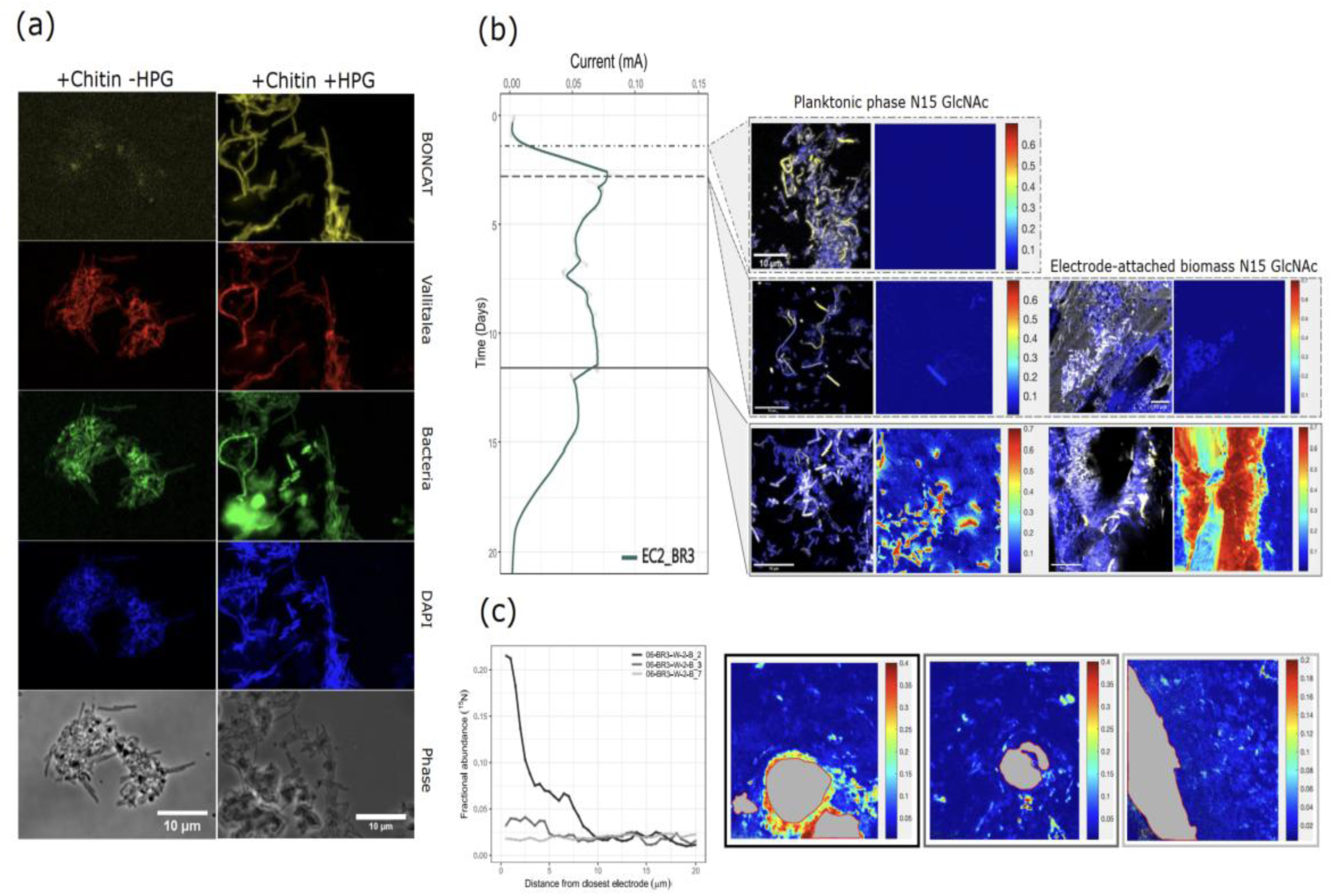
Imaging of planktonic and electrode-associated biomass from the second electrochemical experiment (EC2_BR3). (a) Anaerobic incubations of planktonic biomass amended with fresh chitin and HPG were used to assess translational activity by BONCAT within 5 days of incubation. BONCAT-FISH analysis indicates that *Vallitalea* cells were active with chitin, as the sole carbon and nitrogen source, supporting its role as an anoxic chitin degrader. (b) N15-labeled N-acetyl glucosamine (GlcNAc) was added to EC2_BR3 to examine interactions between *Vallitalea* (planktonic and electrode-attached; visualized with FISH probes depicted in yellow) and spatially proximal microorganisms (DAPI-stained depicted in blue). FISH-nanoSIMS analysis at different current-production stages shows that *Vallitalea* initiated anabolic activity, interpreted through fractional abundance of ^15^N, during the stationary phase of anodic current (μA), followed by increased activity of neighbouring electrogenic cells (metal reducers) and other microbes. (c) Spatial heterogeneity in ^15^N enrichment reveals that the secondary consumers’ anabolic activity was highest in proximity to *Vallitalea* cells and the poised electrode, with activity diminishing beyond 10 µm from the electrode surface (gray shaded region). Hare, three separate NanoSIMS acquisitions were analysed from EC2_BR3 on day 12 of N15-labeled GlcNAc incubation. NanoSIMS image acquisitions were performed with 512 × 512 pixels in a raster size of 35 µm square area.

Previous studies on monospecies *Geobacter sulfurreducens* biofilms respiring on acetate with an electrode showed that metabolic activity in the biofilm decreases significantly at distances greater than 10 µm from the graphite electrode surface^48,49^. To investigate whether similar limitations exist in multispecies GlcNAc degrading EET biofilm, we quantified ^15^N cellular enrichment with distance to the electrode using FISH-nanoSIMS. Our findings with the multi-species microbial biofilm revealed a more heterogeneous response, with variations in average cellular ^15^N enrichment observed within the electrode-attached biofilm community. The ^15^N fractional abundance of the electrode-attached community (BR3, day 12) showed a marked spatial gradient, with average ^15^N value of 20 atom% for cells at the electrode surface decreasing to 2.5 atom% by 10 µm from the electrode. In a separate transect, the ^15^N enrichment dropped from approximately 4 atom% to 2.5 atom% at 5 µm distance (Figure 5c). These gradients illustrate the inherent heterogeneity in metabolic activity within the mixed biofilm community, likely driven by spatial differences in substrate availability and microbial interactions. Unlike polished graphite electrodes, which offer a uniform surface ideal for monoculture studies, the porous carbon cloth electrodes used here provided an expanded surface area and promoted the formation of localized microenvironments which in turn support higher EET activity^50^. Moreover, in our system, metabolic activity depended not only on the availability of the electrode as an electron sink, but also on local substrate production and cross-feeding between GlcNAc degraders with their associated metabolic partner. These single cell activity results affirm the influence of spatial proximity on microbial metabolism in mixed community EET biofilms on poised electrodes.

### Co-cultured deep-sea whale fall isolates, Vallitalea and Trichloromonas, recapitulate anaerobic chitin degradation to EET

The predominance of *Vallitalea* as a primary degrader alongside predicted electrogenic *Trichloromoma*s in the electrode biofilm, opens the question of the minimal number of community members required for anaerobic chitin degradation with EET. We leveraged our bioelectrochemical reactors enrichments to attempt to isolate key members using traditional anaerobic cultivation techniques (see methods). Isolates of anaerobic chitin degraders and metal reducers were successfully recovered from the bioelectrochemical reactor EC2_BR3.

Among these, an isolated strain belonging to the genus *Vallitalea* sp. (sp2), a firmicute, demonstrated enhanced chitinase activity when incubated with 0.2 g of chitin under anoxic conditions in Hungate tubes within 6 days of chitin incubation at 22 °C (Figure 6a). This activity increased progressively with a longer incubation period, indicating the strain’s capacity to degrade chitin over time. Notably, distinct trends were observed in the activity of specific enzymes involved in chitin degradation with this isolate. N-acetylglucosaminidase exhibited the highest activity during the early incubation phase, suggesting a role in the initial breakdown of chitin into smaller polymers or monomeric units. By contrast, endochitinases and chitobiosidases showed increased activity at later stages of incubation. This shift in enzymatic activity over time may be attributed to the antagonistic effects of accumulated byproducts, particularly *N-acetylglucosamine* (GlcNAc), which could inhibit chitinase gene expression^10^.

**Figure 6.**
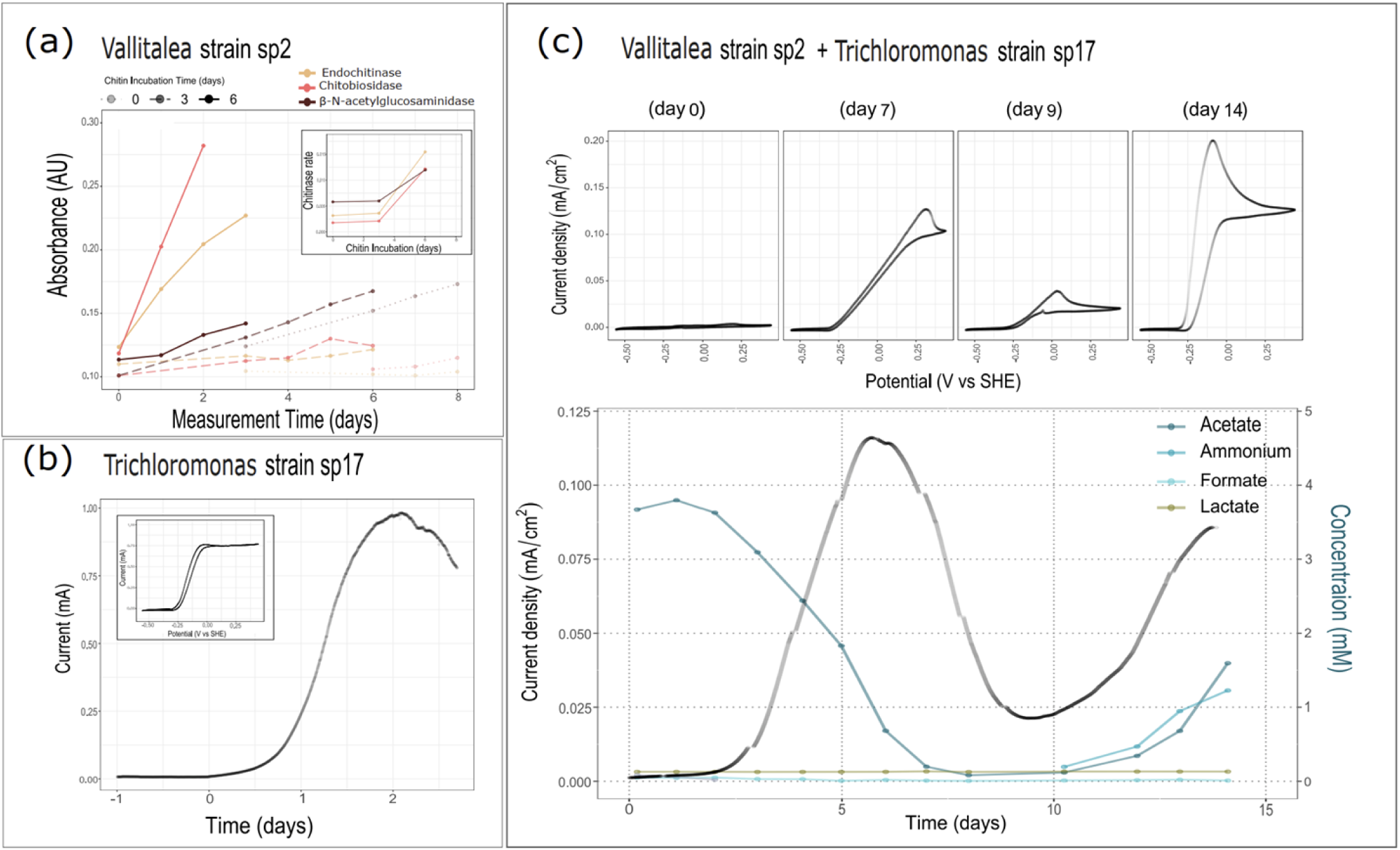
Co-culture of isolates, from EC2_BR1, are capable of anoxic chitin degradation with EET. Investigation of microbial isolates *Vallitalea* sp. (str2) and *Trichloromonas* sp. (str17) (*Desulfuromondaceae*) for anoxic chitin degradation and electrochemical activity. (a) Vallitalea sp. (str2) degrades chitin as evidenced by increased extracellular chitinase activity after 3 days of incubation with chitin. Meanwhile, *Trichloromonas* sp. 17, with 20 mM sodium acetate, uses a poised electrode at +0.22 V vs SHE (carbon cloth) as a terminal electron acceptor for growth. (c) A crossfeeding interaction between *Vallitalea* sp. 2 and *Trichloromonas* sp. 17 was demonstrated in a bioelectrochemical reactor, where the degradation of 0.2 g insoluble chitin was coupled with electron transfer to a poised graphite electrode (surface area: 3.1 cm^2^) serving as an insoluble electron sink. (Bottom) Chronoamperometry data for *Vallitalea* sp. (str2) and *Trichloromonas* sp. (str17) co-incubated shows trends in key metabolites, including acetate and ammonia. Acetate and ammonia concentrations increased by the day 10 of chitin incubation, with an initial acetate decrease attributed to residual acetate from the inoculum. Sulfate (not shown), formate, and lactate remained below 0.2 mM concentration. (Top) A low scan rate (1 mV/s) cyclic voltammogram displays a sigmoidal curve, with an onset potential of-0.25 V vs SHE (similar to EC2 in Figure 3c), indicative of catalytic carbon substrate turnover as both the biomass matures.

Another isolate belonged to the genus *Trichloromonas* sp. (sp17), a *Desulfobacterota*, isolated under anoxic conditions in Hungate tubes with acetate and fumarate amendments at 22 °C (see Methods). *Trichloromonas* sp. (sp17) catalytically oxidized 10 mM acetate using an electrode (carbon cloth) poised at a potential of 0.22 V versus SHE to produce maximum current of 1 mA (Figure 6b). Cyclic Voltammetry (CV) analysis revealed a catalytic turnover onset potential of approximately-0.25 V versus SHE, consistent with previously observed catalytic behavior (EC1 and EC2). However, the CV profile showed notable differences compared to earlier experiments, particularly the absence of additional signature peaks with midpoint potentials of-0.125 V and +0.185 V vs SHE (observed in Figures 2b and 3c). We attribute this difference to the monospecies *Trichloromonas* sp17 biofilm relative to the mixed community in the EC reactors which likely influenced the electrochemical behavior. By minimizing the potential for secondary interactions or competing processes from other community members, this experimental EET setup confirmed and characterised the electrogenic behaviour of *Trichloromonas* sp. (sp17).

Leveraging these isolates, we tested whether a simplified two-member association could successfully couple anaerobic chitin degradation to EET. *Vallitalea sp. (sp2)* and *Trichloromonas* sp. (sp17) were both introduced into an electrochemical reactor equipped with a graphite electrode (3.1 cm^2^) and supplemented with 0.2 g of chitin as a combined carbon and nitrogen source. Initially, the electrochemical reactor was supplemented with 4 mM sodium acetate to facilitate the growth of *Trichloromonas* sp. (sp17) and confirm EET to the poised electrode (Figures 2c, incubation days 0-8). Once the acetate was consumed, an increase in both acetate and ammonia was observed, reflective of active chitin degradation (Figures 2c, incubation days > 8). This increase likely resulted from enhanced substrate turnover, as the growing microbial assemblage metabolized chitin more effectively. Active cross-feeding of metabolic byproducts from chitin produced by *Vallitalea sp. (sp2)* (e.g., acetate), would in turn stimulate the activity and growth of the electrogenic *Trichloromonas* sp. (sp17), further amplifying the current output. This controlled co-incubation experiment demonstrated the synergistic metabolic relationship between these strains, with *Vallitalea sp. (sp2)* acting as a primary degrader and *Trichloromonas sp. (sp17)* acting as a metal reducer and an electroactive organism. This co-incubation provided a direct demonstration of a minimal deep-sea community capable of cooperatively degrading chitin anaerobically and transferring electrons to a poised electrode. By linking insoluble biopolymer breakdown to EET in a defined two-member association derived from deep-sea sediments, we developed a tractable model system to conduct detailed investigations of community assembly dynamics for anaerobic chitin degradation in marine environments and the underlying metabolic pathways required.

## Discussion

In the Monterey Submarine Canyon, whale carcasses (sunken whale fall sites) contribute significant organic material to the seafloor^51,52^ and deep-sea sediments in localized areas (Supplementary Figure 7), offering a unique opportunity to investigate microbial pathways involved in chitin degradation and their coupling to EET processes, which have remained largely unresolved in anoxic deep-sea sediments. As the Whale Fall 1018 (WF1018) site lies in the oxygen minimum zone^30^, it contributes to anaerobic microbial metabolism^34,53^, especially as we observed higher concentration of ferrous iron in the background sediments (ca. 0.14 mM).

Observations from December 2019 (Supplementary Figure 7) show that whale bones persist nearly 20 years post-emplacement, supporting continued local biodiversity and associated enrichment in organic matter, including chitin, to the deep-sea sediments. Our innovative, long-term electrochemical incubations allowed the enriching of microbial communities dominated by anaerobic chitin degraders and iron reducers. The use of bioelectrochemical systems proved instrumental to simulate sedimentary redox conditions (e.g. iron oxide reduction), and we were therefore able to combine the real-time monitoring of electron flow with the manipulation of environmental conditions (C-source) to study microorganisms and metabolic processes of interest (anaerobic chitin degradation coupled to iron reduction; Figure 2a-c). Moreover, our work highlights the effectiveness of bioelectrochemical reactors not only as platforms for the enrichment of EET-capable microbes, but also to electrochemically investigate microbial degradation of insoluble biopolymers, like chitin, by providing sensitive, real-time proxies for detecting the flux of key metabolic intermediates such as acetate (Figure 3d-f).

Building on this, co-occurrence network analysis of the electrode-attached biofilm offered insights on how a stable, functionally cohesive microbial community can emerge from a more complex sediment-derived inoculum through metabolic specialization and redundancy (Figure 4c). Despite a reduction in community complexity compared to the source sediment, the chitin-degrading electrode-respiring communities retained substantial functional diversity. Anaerobic degradation of complex or insoluble polymers like chitin typically requires multiple sequential metabolic steps, often distributed across distinct microbial lineages. Potential coordinated metabolic interactions were exemplified by the co-occurrence of the primary chitin degrader *Vallitalea*, with fermenters and electroactive microbes like *Trichloromonas, Desulfuromonadales,* and *Shewanella*. Several subnetworks emerged from the analysis, with apparent functional and metabolic redundancy among the enriched deep-sea community, including one composed of unknown members within the family *Lachnospiraceae*, *Desulfuromonadaceae* closely connected with an *Abyssivirga*. While another subnetwork grouped *Shewanella*, Clostridium_sensu_stricto_7, and *Sphaerochaeta* together. Notably, the methylotrophic methanogen *Methanolobus* was an unexpected member of the chitin degrading EET community and likely relied on substrates produced by metal reducers, potentially in syntrophic interactions. The large distribution of *Pseudomonas* and *Vallitalea* (on chitin, in the planktonic phase, and attached to the electrode) reflected their metabolic versatility and importance in the stable chitin-degrading community. Ultimately, the electrochemical environment selected for a functionally partitioned community, chitin degraders, fermenters, and EET-capable organisms, demonstrating how selective pressures and metabolic redundancy drive the emergence of efficient, cooperative networks in anoxic systems.

The experimental confirmation of syntrophic interactions, with the co-culture of *Vallitalea sp*. (sp2) and *Trichloromonas sp.* (sp17) demonstrated the importance of substrate sharing and interspecies cooperation in anoxic chitin degradation. We were able to show that the *Firmicute* V*allitalea sp. (sp2)* acts as a primary degrader of chitin, breaking down the insoluble biopolymer into fermentative byproducts such as acetate. These metabolites sustained the activity of secondary consumers, including *Trichloromonas sp. (*sp17), closely related to the well-characterized iron-reducing bacteria *Desulfuromonas*^37,54^. While previous studies have identified *Vallitalea* as a putative chitin degrader^41^ and *Trichloromonas* as putatively capable of EET^55^, their cooperative interaction in the context of deep-sea polymer degradation had never been studied. Our results therefore provide the demonstration of a cooperative partnership enabling chitin degradation coupled to iron oxide respiration via EET. This division of labor sustained metabolic activity, enabling effective redox coupling between chitin and iron oxides, two insoluble substrates. The metabolic activity within this biofilm community evolved in time, with *Vallitalea* (planktonic phase and electrode-attached) becoming active first, followed by electroactive microbial neighbors, presumably *Trichloromonas* (predominantly electrode-attached). Notably, the electroactive microbes, *Trichloromonas*, were more metabolically active in the chitin fed reactor when both their metabolic partner (*Vallitalea*) and the electrode were in close spatial proximity (Figure 5c). Moreover, the overall anodic activity of the electrode-attached mixed species biofilm declined beyond 10 µm distance from the electrode surface. These findings support the idea that “metabolism at a distance” imposes energetic costs that may inhibit the growth and activity of electrode-attached microbial communities^48,49^, emphasizing the requirements for a specific spatial organization to enable metabolic cooperation in such systems.

By functionally linking chitin degradation to metal oxide respiration, this work reveals how the microbial communities in organic rich deep-sea sediments contribute to organic matter turnover and the mobilization of bioavailable nutrients. This mechanism is particularly relevant in oxygen-limited deep-sea environments, where alternative electron acceptors play a vital role in chitin degradation, sustaining microbial activity and ecosystem functionality^56^. Related studies found that chitin is degraded and metabolised in both aerobic environments such as soil^57,58^, marine water column^12^, marine sponges^59^, and anaerobic freshwater sediments^60^. However, none of these investigations have explored spatial challenges of coupling degradation of an insoluble chitin polymer to metal oxide respiration through EET. Our research addresses this gap by elucidating the mechanisms and microorganisms involved in anoxic chitin degradation from deep-sea sediments. While the spatially segregated microbial communities in the bioelectrochemical reactors were complex, our results demonstrate that successful chitin degradation coupled to EET is possible with a simple two-member system, opening new possibilities as a defined, tractable model for ecophysiological studies. Further detailed investigations about the interspecies interactions that drive nutrient cycling and redox balance in anoxic marine ecosystems will deepen our understanding of the complex metabolic networks sustaining these environments. Overall, our research offers a framework for leveraging bioelectrochemical systems to study community assembly associated with complex anaerobic carbon cycling in anoxic environments.

## Methods

### Whalefall site characteristics and sediment collection

Whale Fall 1018 (called WF1018) was a 17 m long blue whale that was implanted in the Monterey submarine canyon (36.771442 N, 122.082998 W) in October of 2004. This whale fall is located in the oxygen minimum zone, defined by oxygen values below 0.5 mg/L^30^. In Dec 2018, during a research cruise WF12-18 on the R/V Western Flyer, marine sediment core was collected by the ROV Doc Ricketts from this former whale fall site. The background core, located less than 2 m distance, showed a spike in porewater iron (0.13 mM Fe^2+^) just below the sediment-water interface at a depth of 1–2 cm. Sediment cores were extruded upwards, sliced into 1 cm horizons, and stored at 4 °C in an argon-gassed Mylar bag.

### Iron enrichment

Sediments recovered from the 1-2 cm depth horizon were used in our iron oxide culture enrichments. Chitin and poorly crystalline iron oxide (PCIO) enrichment cultures were prepared in 60 mL serum vial containing 20 mL modified Artificial Sea Water (ASW) media containing (in g/L): NaCl, 24; MgCl _2_.6H2O, 10.67; CaCl_2_.2H_2_O, 1.3; KCl, 0.67; KBr, 0.1; H_2_BO_3_, 0.027; SrCl_2_.6H2O, 0.027; NaF, 0.003; Na_2_SO_4_, 1.5. Chitin (0.01 g/mL) was added as carbon and nitrogen source, and poorly crystalline iron oxide (PCIO), as the terminal electron acceptor (0.013 g/mL).

HEPES buffer (25 mM) was used to maintain the pH at 8.2. Trace elements and vitamins used were SL-11 and Modified Wolin’s mineral solution DSM141, respectively, as provided by Media*Dive* (DSMZ)^61^. PCIO mineral was prepared by adapting a previous recipe^62^. We added 12 g of FeCl_3_.6H_2_O in 200 mL of milliQ water (final concentration of 0.4 M). The mixture was constantly stirred and slowly adjusted pH to 7.0 by adding 10 M NaOH dropwise. To remove dissolved chloride, centrifuge the suspension at 5000 RPM for 15 min (repeat 6 times). The final pellet was freeze-dried (or speed vacuumed at 60 °C at maximum speed overnight) and stored as fine powder at-80 °C. The cultures were incubated at 10 °C and 22 °C under anoxic conditions. Enrichment of an active iron-reducing microbial community was determined by ferrozine colorimetric assay^63^.

### Electrochemical enrichment

The chitin-PCIO-enriched was inoculated into triplicate bioelectrochemical reactors operated first at 10 °C for 35 days, followed by 22 °C. A standard three-electrode glass cell comprised of (1) a working electrode (WE) as a carbon cloth with dimensions 1 × 1 cm, (2) a counter electrode (CE) as a platinum wire (CHI115, CH Instruments, USA), and (3) a reference electrode (RE) as a 1 M KCl Ag/AgCl reference electrode (CHI111, CH Instruments, USA). Working potential and current produced were maintained at +0.22 V vs SHE and constantly monitored, using a potentiostat (Squistat Prime, Admiral Instruments, USA). The electrochemical reactor carried a total volume of 40 mL, and its headspace was constantly purged with N2 gas to maintain an anoxic environment. The planktonic phase of reactors was continuously adjusted for any change in salinity and pH. Cyclic voltammetry was performed at the scan rate of 1 mV/s for 3 cycles (See Supplementary Materials for detailed CA and CV profile for echem_run1, EC1 and echem_run2, EC2).

### Ion chromatography and Metabolomics

The planktonic phase of each reactor (2 mL) were sampled (Supplementary Materials) and then filtered using 0.2 µm PES filters. The filtered samples were then stored at-20°C for subsequent analysis, including ion chromatography (IC) and exometabolomics. For IC, samples were diluted 1:50 in Type 1 ultrapure water and run on Dionex ICS-2000 or Thermo Integrion HPIC as previously described^64^ using an AS19 anions and CS16 cations column. Samples were run at the Water and Environment Laboratory (Caltech). Exometabolomics samples were sent to ETH, Zurich for further analysis. Untargeted metabolite profiling and analysis were conducted using Flow Injection Analysis Quadroupole Time of Flight Mass Spectrometry (FIA-QTOF-MS). To prepare the samples, all supernatants were diluted 100-fold in water before measurements. Metabolomics analysis was performed using a binary LC pump (Agilent Technologies) and an MPS2 Autosampler (Gerstel), coupled to an Agilent 6520 time-of-flight mass spectrometer (Agilent Technologies). The mass spectrometer was operated in negative mode, 2 GHz, extended dynamic range, with a mass range of 50-1000. The mobile phase consisted of a mixture of isopropanol and water (60:40, v/v), supplemented with a 5 mM ammonium fluoride buffer at pH 9, and the flow rate was set to 150 μl/minute. All raw data obtained from the measurements underwent spectral processing and alignment using Matlab software (The Mathworks, Natick), following established procedures^65^. In total, we identified 2976 ions, out of which 158 ions were annotated based on exact mass. The annotation process involved comparing the ions against a curated compound library of metabolites predicted to be present in taxa found in our environmental samples, using BioCyc metabolic networks as a reference^66^. A tolerance of 0.005 Da was applied during the annotation process. In cases where a single m/z ion matched multiple compounds with isomeric or isobaric characteristics in the compound library, the top annotation was assigned to the compound that participates in the largest number of enzymatic reactions across the pathway databases. To identify metabolites in each bioreactor, we applied a cutoff requiring ion intensities to exceed two-fold over those in the blank medium. See Supplementary Materials for detailed IC and exometabolites profile for echem_run1, EC1 and echem_run2, EC2.

### Exoenxyme assay for chitin degradation

Chitinase activity within the planktonic phase of the bioelectrochemical reactors (EC2_BR1 and EC2_BR3) were monitored over a 40-day chitin incubation period. Enzymatic hydrolysis of chitinase substrates, indicative of β-N-acetylglucosaminidase, chitobiosidase, and endochitinase activity, was quantified by measuring the release of p-nitrophenol using a colorimetric assay at 405 nm (Kit CS0980, Sigma-Aldrich). To perform the assay, 500 µL of the planktonic phase were aseptically sampled and filtered through a 0.2 µm sterile PES filter. A 195 µL volume of the filtered sample was incubated with 5 µL of each of the three substrates, with reactions conducted individually for each substrate in replicates. Colorimetric measurements were taken daily to monitor the progress of enzymatic activity over time. The rate of chitinase activity was plotted to illustrate the temporal dynamics and evolution of enzymatic activity in the chitin-fed reactor.

### Nucleic acid extraction

High molecular weight DNA extraction was performed from various samples (see Supplementary Materials) using phenol-chloroform extraction. Briefly, samples were resuspended (chitin-associated biomass and electrode-attached biomass) or added (0.5 mL of planktonic phase) in 0.5 mL of freshly prepared filter sterilized sucrose lysis buffer (40 mM EDTA, 50 mM Tris HCL, 0.75 M sucrose) and mixed by inverting. Cell lysis was performed by incubating with lysozyme (2 mg) at 37 °C for 45 min, followed by protein digestion by incubating with proteinase K (40 U) and 2% of sodium dodecyl sulfate (SDS) solution at 37 °C for 30 min. For higher throughput, incubation duration in proteinase K and SDS was increased to 1.5 hours at 60 °C. Finally, equal amounts of phenol:chloroform:IAA (25:24:1) at pH 8 were added to the homogenized mixture and mixed by vortexing for 30-60 sec. The organic and aqueous phases were separated by centrifugation for 5 min at 16000 x g. The aqueous phase containing the genomic DNA was then collected, and extracted in phenol:chloroform:IAA solution to increase purity of the extracted nucleic acids. Finally, the remaining phenol was removed by adding an equal amount of chloroform:IAA (24:1), vortexing (30-60 sec), and centrifuging at 16000xg for 5 min. The genomic DNA was concentrated using ethanol precipitation. Here, sodium acetate (0.3M) was added, followed by 100% ethanol to a final volume four times that of the aqueous phase. This mixture was incubated overnight at-20 °C. The next day, DNA was precipitated by centrifugation for 20 mins at maximum speed at 4 °C. The ethanol supernatant was carefully removed without disturbing the DNA pellet. The pellet was then resuspended in 10 mM Tris and any remaining ethanol was removed using multiscreen filter plates and vacuum manifold following the manufacturer’s instruction (Merck Millipore, USA). The extracted DNA was resuspended in 30-50 mL of 10 mM Tris (pH 8) and quantified using a UV spectrophotometer or Qubit fluorometer dsDNA assay. Extracted DNA was stored in-80 °C until further downstream analysis.

### 16S rRNA gene amplicon sequencing

The V4-V5 region of the 16S rRNA gene was amplified using 515F and 926R [Parada, 2016] archaeal/bacterial primers with Illumina adapters (515F 5’-TCGTCGGCAGCGTCAGATGTGTATAAGAGACAG-GTGYCAGCMGCCGCGGTAA-3’; 926R 5’-GTCTCGTGGGCTCGGAGATGTGTATAAGAGACAG-CCGYCAATTYMTTTRAGTTT-3’). PCR reactions were done in duplicate with Q5 Hot Start High-Fidelity 2x Master Mix (New England Biolabs, MA, USA) in 15 μL reaction volumes according to manufacturer’s directions with annealing conditions of 54°C for 30-35 cycles. Duplicate PCR samples were pooled and barcoded with Illumina Nextera XT index 2 primers that include unique 8-bp barcodes using Q5 Hot Start with 3 μL duplicate-pooled PCR product in a 30 μL reaction volume, annealed at 66°C, and cycled 11 times. Products were run on 1.5% agarose gel and quantified by band intensity. Barcoded PCR products were combined in equimolar amounts and 300uL of this combined sample was run on 1.5% low melt agarose gel and purified with Promega’s Wizard SV Gel and PCR Clean-up System (Madison, WI, USA). The sample was sequenced by Laragen (Culver City, CA, USA) using the MiSeq Reagent Kit v3 (600-cycle) #MS-102-3003 on Illumina’s MiSeq platform with the addition of 15-20% PhiX. The samples sent for sequencing and analysed downstream are provided in Supplementary_table_16S_metadata.xlsx and the raw data is uploaded on NCBI.

### 16S rNA gene sequence analysis

Sequence data were processed in the 2020 distribution of QIIME2 software suite (Qiime2-2020.11)^67^ and the metadata were validated using the browser supported tool, Keemei, to check for any issues with metadata^68^. Primer sequences were removed using Cutadapt^69^. The trimmed sequences were then loaded into QIIME2, followed by merging and denoising steps via DADA2^70^. The taxonomy classifier was trained on the SILVA 138 database using RESCRIPt^71^. Contaminants were removed sequentially 1) ASV’s present in < 2 samples, 2) singleton ASVs, 3) ASVs unique to control samples, 4) potential contaminants were statistically inferred through decontam^72^ and manually removed. Finally, all the control samples were removed to generate 140 samples and 1568 ASVs remaining as per ASV table. A phylogenetic tree was built using MAFFT^73^. The data were then exported for further analysis in R.

Alpha diversity (Chao1 and Shannon’s diversity indices) and beta diversity comparisons (NMDS with Bray-Curtis dissimilarity, stress: 0.145) were evaluated for each incubation using the Phyloseq package^74^. Each data point represents a sample taken at various sediment depths (from whale fall sediments) and at different time points across iron and electrochemical incubations.

To assess alpha diversity, the mean index values for whale fall sediments and corresponding enrichment samples were calculated and compared using a Wilcox signed-rank test, with p-values adjusted using the Bonferroni correction method to determine significant differences using rstatix^75^. Beta diversity was visualized by plotting microbial community shifts across incubation types, using the mean abundance of each ASV within the same incubation. A stream plot was employed to visualize the evolution of microbial community composition through subsequent enrichment stages. To identify the taxa contributing most to dissimilarity between sample types, SIMPER (Similarity Percentage) analysis was performed^76^. Additionally, indicator species analysis (ISA)^77^ using the indicspecies package^78^ identified key taxa responsible for variations in microbial community structure between whale fall sediments and laboratory incubations.

Differential abundance analysis was conducted using ANCOM-BC^79^ to evaluate changes in microbial abundance across different phases of the electrochemical reactor (electrode-attached, planktonic, and chitin-attached). The analysis employed false discovery rate (FDR) correction for p-value adjustments. Absolute abundances of the resulting differentially abundant taxa were visualized using jitter box plots (Figure 2e), alluvial diagrams (Figure 4a), and stream plots (Figure 4b).

Microbial associations were analyzed and visualized as a network using the NetCoMi ^80^ package, applying Pearson correlation and CLR (Centered Log-Ratio) normalization. Network clustering was performed using the fast greedy algorithm, and the network was visualized using the iGraph tool. The full data analysis pipeline, including the R Markdown scripts, is provided as a Supplementary Materials (16S_iTag_analysis_chitin_echem.html).

### Fluorescence in situ hybridization (FISH) probe design (SAC)

A new FISH probe was designed using the ARB 6.0.6 software program^81^ and the Silva database SILVA_138_SSURef_NR99_05_01_20_opt^82,83^. It was designed to specifically target *Vallitalea* sp with the sequence (5’-CGACCCCCGACACCTAGCAT-3’).

### Stable Isotope Probing

Stable isotope probing was conducted using biological replicate 2 from the second electrochemical run. The reactor was incubated with 3 mM ¹⁵N-labeled GlcNAc. Heavy isotope-labeled medium was identical in chemical composition to the previously described (see methods Iron enrichment) modified ASW medium, with an increase in the final heavy isotope of 15N to 98 atom% in GlcNAc. Enriched isotopic chemicals were purchased from Cambridge Isotope (NLM-8810-0.1). The isotopically labelled ^15^N-GlcNAc (3mM) was directly added to the bioelectrochemical reactors without any prior medium exchange. Anodic current was continuously measured (Figure 5b). Planktonic phase samples were collected at days 1.5, 3, and 12. Additionally, small sections of the poised electrode (carbon cloth) were sampled on days 3 and 12 for downstream analysis. All the samples were collected in anaerobic conditions. For negative control samples to test for nonspecific isotope binding, a kill control of ^15^N-GlcNAc incubated planktonic cells (autoclaving at 121 °C for 45 min) was employed (Supplementary Figure 5).

### Sample FIxation

Samples (planktonic phase, chitin and carbon cloth) were fixed by adding paraformaldehyde (PFA) to a final concentration of 2%, followed by incubation at 37 °C for 10 minutes or at 4 °C overnight. To quench the unreacted PFA and to protect sample quality, an equal volume of 750 mM Tris-HCl buffer (pH 8) was added, and the samples were incubated at room temperature for 20 minutes. The samples were next washed twice with 1X PBS to remove residual PFA and subsequently stored in 1X PBS containing 50% ethanol, until further analysis.

The carbon cloth samples were embedded in glycol methacrylate resin (Technovit 8100, Kulzer, Germany) and sliced into thin sections by microtome with thickness of *ca.* 5 µm as previously defined^48^.

### Fluorescent in-situ Hybridization (FISH)

The phylogenetic identity of microorganisms in the samples (planktonic phase, chitin-attached, and electrode-attached) were determined using conventional FISH using oligonucleotide probes fluorescently labeled on both the 5’ and 3’ ends (dual labeled) as outlined below. For first electrochemical run (echem_run1, EC1), the FISH probes used were: 1) general deltaproteobacterial probe DELTA495a (FAM): 5’ to 3’ = AGTTAGCCGGTGCTTCCT^84,85^, 2) general alphaproteobacterial probe ALF968 (cy5): 5’ to 3’ = GGTAAGGTTCTGCGCGTT^86^, 3) general gammaproteobacterial probe GAM42a (Cy3): 5’ to 3’ = GCCTTCCCACATCGTTT^86^, and 4) an unlabelled competitor probe for GAM42a, BET42a: 5’ to 3’= GCCTTCCCACATCGTTT^87^. For electrochemical run (echem_run2, EC2), the FISH probes used were: 1) general archaeal probe ARCH915 (Alexa647), dual labeled: 5’ to 3’= GTGCTCCCCCGCCAATTCCT^88^; 2) *Valitallea* sp. specific probe Aby-Val-645 (Cy3): 5’ to 3’= CTCCTGCACTCTAGCAAAGC (this study); 3) and another *Valitallea* sp. specific probe Aby-Val-823 (Cy3): 5’ to 3’= CGACCCCCGACACCTAGCAT (this study); 4) general deltaproteobacterial probe DELTA495a (Alexa488), 5’ to 3’= AGTTAGCCGGTGCTTCCT^84^; 5) an unlabelled competitor probe for DELTA495a, DELTA495a_comp 5’ to 3’= AGTTAGCCGGTGCTTCTT^89^. FISH hybridization was conducted as previously described^90^. The carbon cloth samples were hybridized in a hybridization buffer containing 35% formamide and incubated at 46 °C for 2 hrs, followed by a wash step at 48 °C for 15 min.

### NanoSIMS

Sectioned slices, after confocal fluorescence imaging, were rinsed by DI water 3 times. After drying, sections were gold sputter coated (20 nm) to enhance conductivity prior to secondary ion mass spectrometry analysis performed on a NanoSIMS 50L instrument (Cameca) in the Caltech Microanalysis Center (CMC). The gold coated samples were pre-sputtered with a 60-pA primary Cs+ ion beam (aperture diaphragm D1 = 2) until the ^14^N^12^C^−^ ion counts stabilized. Data were collected using a 1.5-pA beam (D1-3) for ions (^14^N^12^C^-^, ^15^N^12^C^-^) for the determination of ^13^C/^12^C, and ^15^N/^14^N ratios, respectively. Acquisitions were performed with 512 × 512 pixels in a raster size of 35 µm square area. Roughly 30 min per frame was used and 2-4 frames were collected depending on samples. Look@NanoSIMS Matlab GUI^91^ was first used to export raw data from nanoSIMS.im format files. All the subsequent data processing and analysis used an in-house MATLAB code (Release 2020b). Regions of interest (ROIs) were manually segmented by using ^14^N^12^C^-^ raw frame as a reference. ROIs with clear contours were selected. For each pixel of the electrode-attached biofilm, the shortest distance to the nearest electrode surface was calculated using the pairwise distance measurements. These pixels were then binned in 0.5-μm increments based on their proximity to the electrode. For each bin, the counts of ^15^N^12^C^−^ and ^14^N^12^C^−^ were summed to compute the fractional abundance of the labeled heavy isotopes using: ^15^*F* = ^15^N^12^C^−^/(^15^N^12^C^−^ +^14^N^12^C^−^). This approach was previously described by Chadwick et al^48^.

### BONCAT-FISH

The planktonic phase of the chitin-fed reactor was incubated in nitrogen-purged, chitin-modified artificial seawater (ASW) media supplemented with 50 µM of the alkyne-bearing artificial amino acid homopropargylglycine (HPG). This setup was designed to identify the translationally active member of the chitin-degrading microbial community. After 5 days of incubation at 22 °C, samples were collected, fixed with 2% paraformaldehyde (PFA), washed with 1X PBS, and stored at-20 °C in 70% ethanol:PBS until further processing.

Bio-orthogonal non-canonical amino acid tagging (BONCAT) was performed using AF647-picolyl azide to detect translationally active cells. The fixed planktonic samples were immobilized on Teflon-coated glass slides and dried at 46 °C. Slides were subsequently dehydrated through an ethanol series of increasing concentration (50%, 80%, and 96% v/v in double-distilled water) and then air-dried completely. A freshly prepared “click cocktail” was used to label incorporated non-canonical amino acids. The cocktail consisted of the following components in 0.2-μm-filtered 1× phosphate-buffered saline (PBS, pH 7.4): 5 mM sodium ascorbate, 5 mM aminoguanidine, 100 μM CuSO₄, 0.5 mM THPTA (tris-hydroxypropyltriazolylmethylamine; copper-stabilizing ligand), 20 μM

AF647-picolyl azide. A total of 20 μL of this “Click Cocktail” solution was applied to each sample of the glass slide. The glass slide was incubated for 60 minutes at room temperature in a humid chamber. Following incubation, slides were rinsed thoroughly with double-distilled water to remove unbound reagents. Following BONCAT, ethanol-washed samples were hybridized with oligonucleotide probes. Fluorescence in situ hybridization (FISH) was performed on the labeled samples as described previously (see FISH Methods section), targeting *Vallitalea* with Cy3-labeled probes and the broader bacterial community with Alexa488-labeled EUB338 (5’-GCTGCCTCCCGTAGGAGT-3’), EUB338-II (5’-GCAGCCACCCGTAGGTGT-3’) and EUB338-III (5’-GCTGCCACCCGTAGGTGT-3’) probes, allowing simultaneous visualization of taxonomic identity and translational activity. The protocol for performing the BONCAT-FISH is taken from previous literature^92^. Samples were mounted with 1 mg/mL 4,6-diamidino-2-phenylindole (DAPI; Sigma-Aldrich) in Citifluor AF-1 antifading solution (Electron Microscopy Sciences) and analyzed using Zeiss ELYRA S1 (SR-SIM) super resolution microscope.

### Microbial isolation

Samples collected from both the planktonic phase and the electrode-attached biofilm of the second electrochemical enrichment (echem_run2, BR1) were used for microbial isolation experiments. These samples were incubated in a modified artificial seawater (ASW) medium (see Methods: Iron enrichment) specifically designed to support the growth of microbes with distinct functional traits. Two types of enrichment media were prepared to selectively target different metabolic capabilities. The first medium was supplemented with 3 mM

N-acetylglucosamine (GlcNAc), a monomer of chitin, to promote the growth of chitin-degrading microorganisms. The second medium contained 10 mM sodium acetate and 20 mM sodium fumarate to enrich microorganisms capable of extracellular electron transfer (EET). Fumarate, a well-established soluble electron acceptor, supports anaerobic respiration in model EET organisms such as *Geobacter sulfurreducens*^93^ and *Shewanella oneidensis*^38^.

To obtain pure cultures from these enrichments, microbial isolations were performed using a dilution-to-extinction strategy in separate anaerobic Hungate tubes. Each set of tubes was tailored to enrich either chitin-degrading or mineral-reducing bacteria. This approach enabled the isolation of functionally distinct microbial strains that had been enriched in the bioelectrochemical reactors (EC2_BR1) facilitating downstream phylogenetic and physiological analyses of isolates. Frozen stocks of the isolated culture were prepared using 10% DMSO as a cryoprotectant inside an anaerobic chamber to maintain viability under oxygen-free conditions. Upon re-inoculation into defined media, visible microbial growth was observed within one month, confirming successful preservation and revival of the isolate.

Ribosomal 16S sequences were obtained for the three isolates using direct 16S rRNA amplification from pure culture DNA extracts. The universal bacterial primers 515F (5′-GTGCCAGCMGCCGCGGTAA-3′) and 1492R (5′-GGTTACCTTGTTACGACTT) were used. Approximately 20–40 ng of PCR product from each isolate were extracted and sent for sanger sequencing by Laragen. The 900 bp long sequences were quality checked and assembled using Geneious 7.1^©^ (Biomatters, New Zealand).

Phylogenetic analysis was performed using the SILVA 138.1 reference database within the ARB software environment (version 7.1.0)^81^. The phylogenetic tree was constructed using the Maximum Likelihood algorithm implemented in RAxML (version 8)^94^. The tree was rooted using *Shewanella* and *Vibrio* as outgroup taxa (Supplementary Figure 6).

## Acknowledgements

We are grateful to members of the Orphan lab for discussions regarding this work. Dr. Daniel Utter provided critical readings that improved this manuscript. This research was supported by the SIMONS foundation Life Sciences-Simons Collaboration on Principles of Microbial Ecosystems (PriME); Award XXXXXXX). Ship time on the R/V Western Flyer and deep-sea sampling with the ROV Doc Ricketts was made possible by the Monterey Bay Aquarium Research Institute (MBARI) where VJO is an adjunct investigator. We extend our gratitude to Dr.

Nathan Dalleska for training and access to the Resnick Water and Environment Lab (WEL), Dr. Giada Spigolon for guidance with LSM 980 hosted at Beckman Biological Imaging Center, and Dr. Yunbin Guan for his expertise with nanoSIMS operations. We also thank Dr. Doug LaRowe and Dr. Grayson Boyer for insightful discussions on Gibbs free energy calculations. We thank Emmanuelle Botté (Manuscribe) for her support in reviewing and editing this manuscript. VJO is a CIFAR research fellow in the Earth 4D program.

## Author Contributions

YJ and VJO conceived the work. Field work was performed by SL and VJO, and subsequently YJ performed experiments with help from SL, FW, YG, SC, SP, and JS. Data analysis and interpretation was done by YJ, YG, and SP. Original draft was written by YJ, refined by VJO and EB with contributions from all authors.

## Data and Code Availability

All raw data and analysis code used to generate the figures, extended data, and supplementary materials supporting the statements in this manuscript are available via the Caltech Research Data Repository (Ref: 10.22002/5hw6d-sw55). The raw sequencing data have been uploaded to National Center for Biotechnology Information (NCBI) Sequence Read Archive (SRA) database (accession number: XXXXXXX) under BioProject YYYYYYY.

---

**Supplementary Figure 1:**
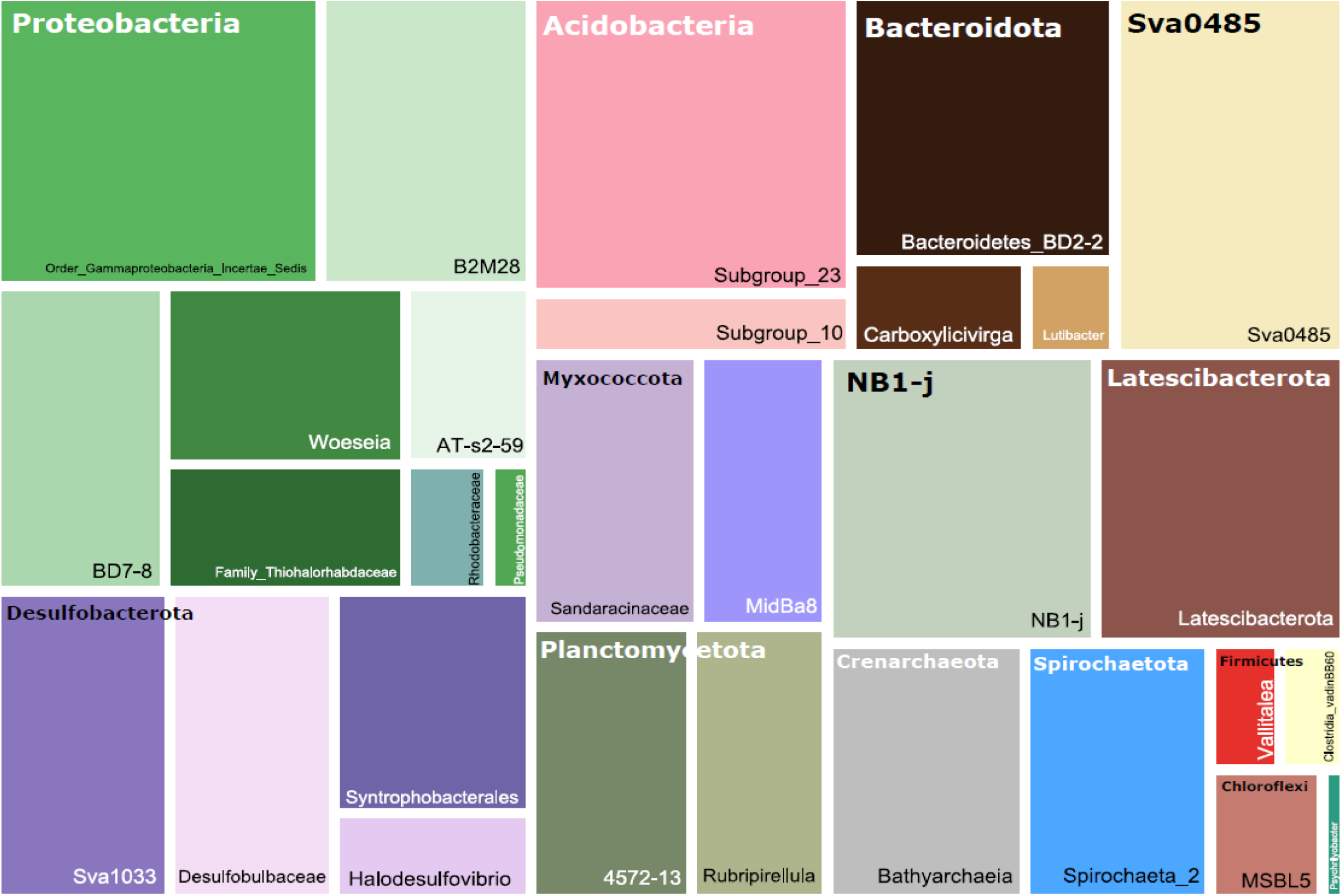
Relative abundances of major archaeal and bacterial phyla identified across sediment samples, highlighting *Proteobacteria, Acidobacteria, Bacteroidota, Sva0485, NB1-j, Latescibacterota, Planctomycetota, Crenarchaeota, Spirochaetota, Firmicutes, Chloroflexi, Myxococcota,* and *Desulfobacterota*. These phyla represent the diverse microbial community structure observed in the sediment core samples, with dominant lineages from *Desulfobacterota* and *Proteobacteria*, and minor representation from phyla such as *Chloroflexi* and *Firmicutes*.

**Supplementary Figure 2:**
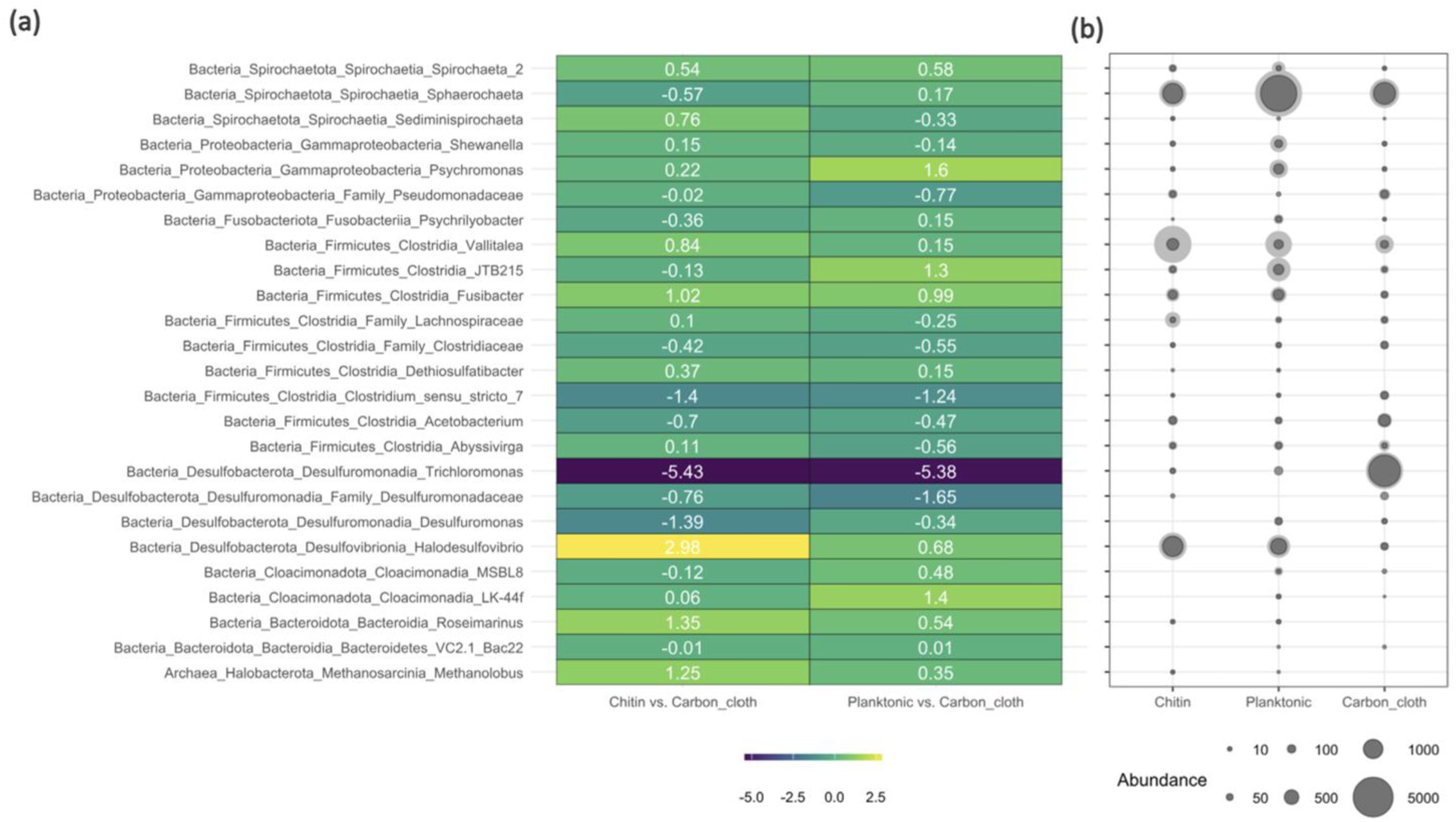
(a) ANCOM-BC heatmap of natural log fold changes in 16S rRNA gene relative abundance for pairwise comparisons (adjusted p < 0.05) with chitin versus carbon cloth (electrode-associated) and planktonic versus carbon cloth (electrode-associated) microbial community for BR3 in echem run 1 (EC1_BR3). (b) Bubble plot showing relative abundances of the respective microbial taxa across chitin, planktonic, and carbon cloth phase.

**Supplementary Figure 3:**
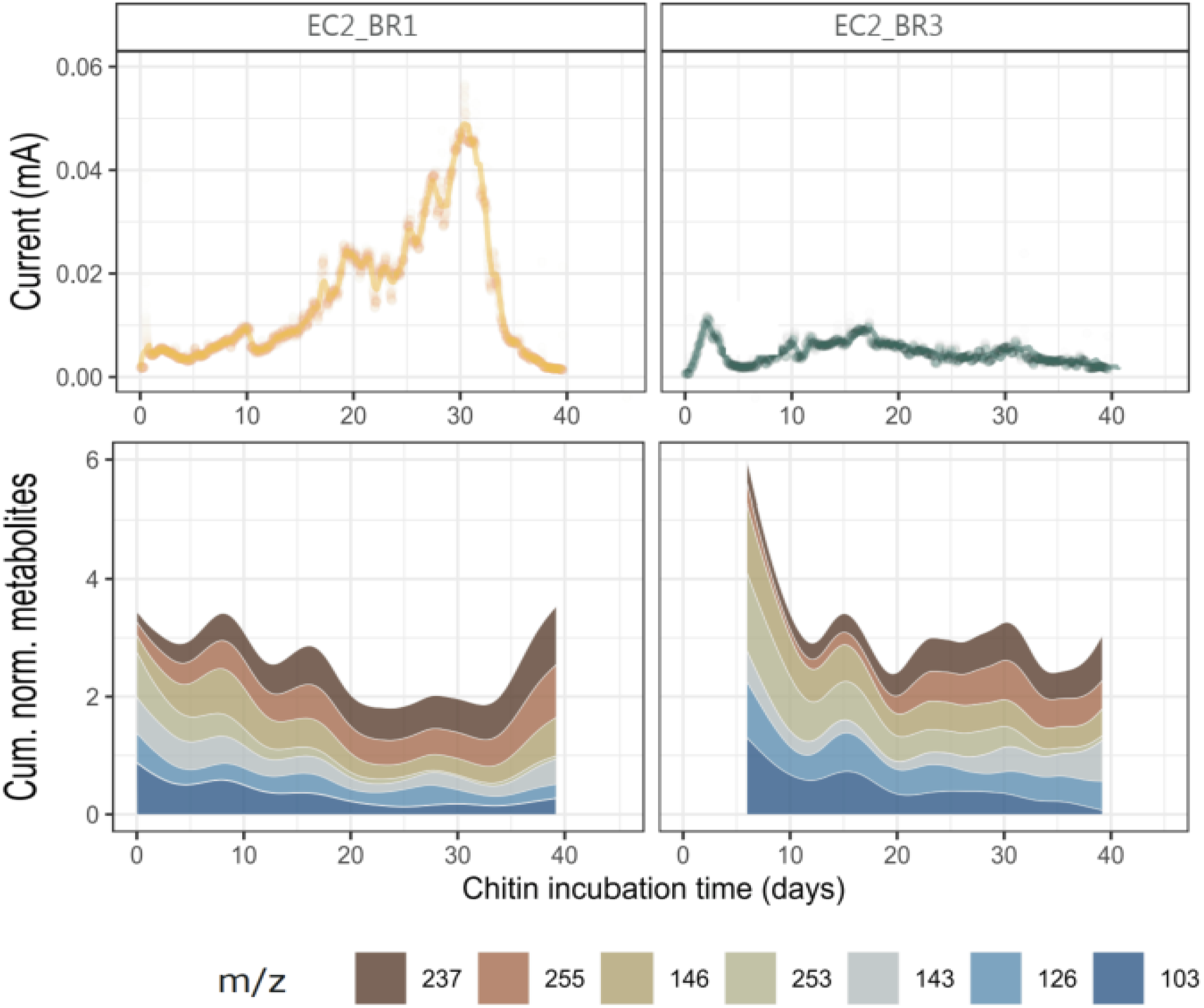
Exometabolites analysis measured in chitin incubation of EC2_BR1 and EC2_BR3 shows trends of 7 unannotated metabolites (with m/z) provided as below. Samples were collected daily.

**Supplementary Figure 4:**
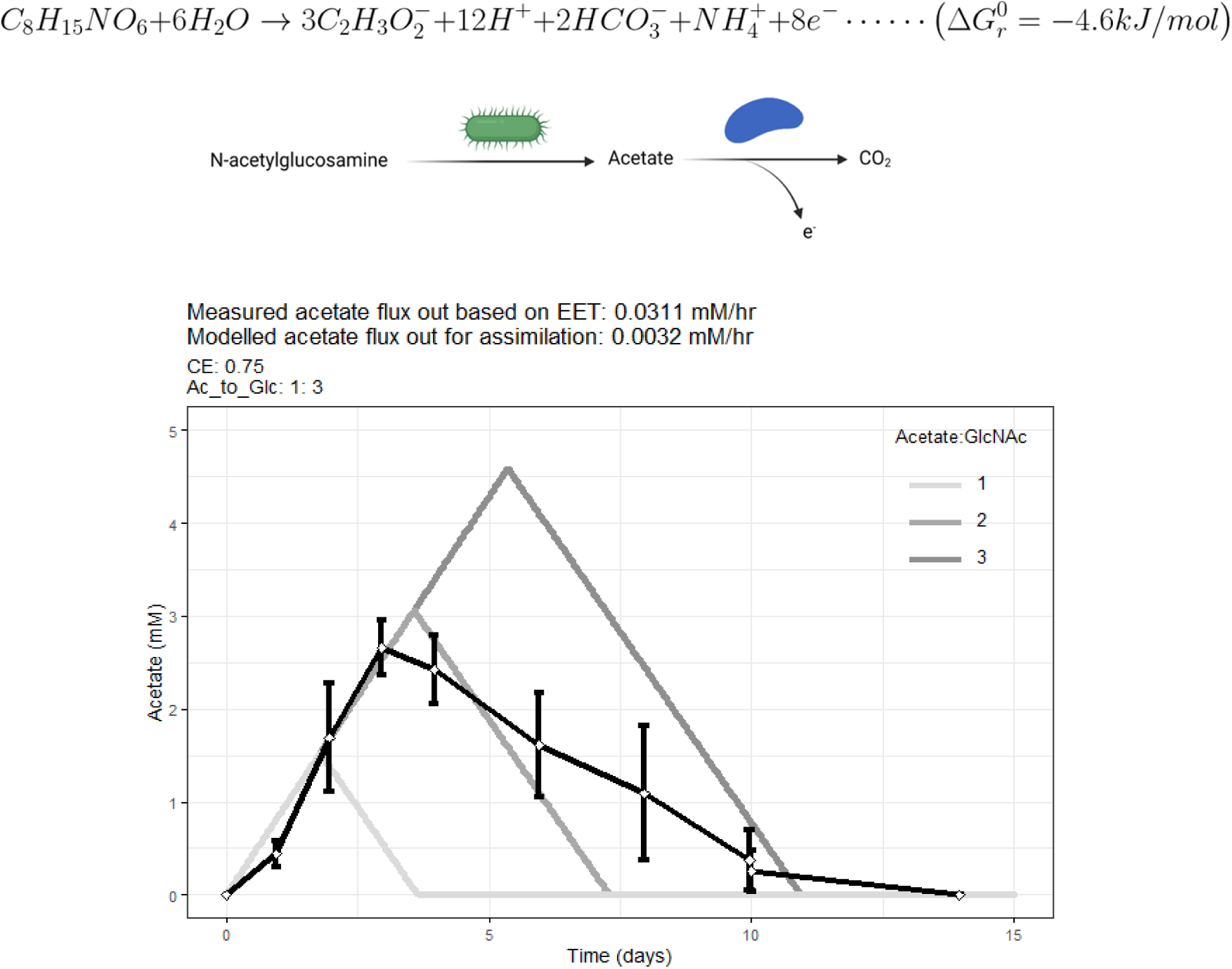
Mathematical model depiction of the conversion of GlcNAc into acetate through fermentation, with acetate serving dual roles: as a substrate for oxidation via EET and as a carbon source for biomass assimilation. The standard Gibbs free energy change for GlcNAc fermentation coupled to acetate oxidation is −4.56 kJ/mol, with each GlcNAc molecule potentially yielding 1, 2, or 3 acetate molecules, one ammonia molecule, and eight terminal electrons. The mathematical model simulates acetate profiles under different acetate:GlcNAc ratios (1:1, 2:1, and 3:1), shown as varying gray lines by using experimentally derived parameters: a GlcNAc degradation rate of 0.07 mM/hr and an electron production rate of 0.0311 mM/hr (based on average anodic current of 200 μA), and acetate assimilation rate of 0.0032 mM/hr. The experimental acetate concentration (black curve) suggests the ratio lies between 1 to 3, due to the heterogeneity within the microbial population. This supports the hypothesis that acetate released during GlcNAc metabolism is sufficient to sustain the activity of EET-capable metabolic partners. The model assumes a coulombic efficiency of 75%.

**Supplementary Figure 5:**
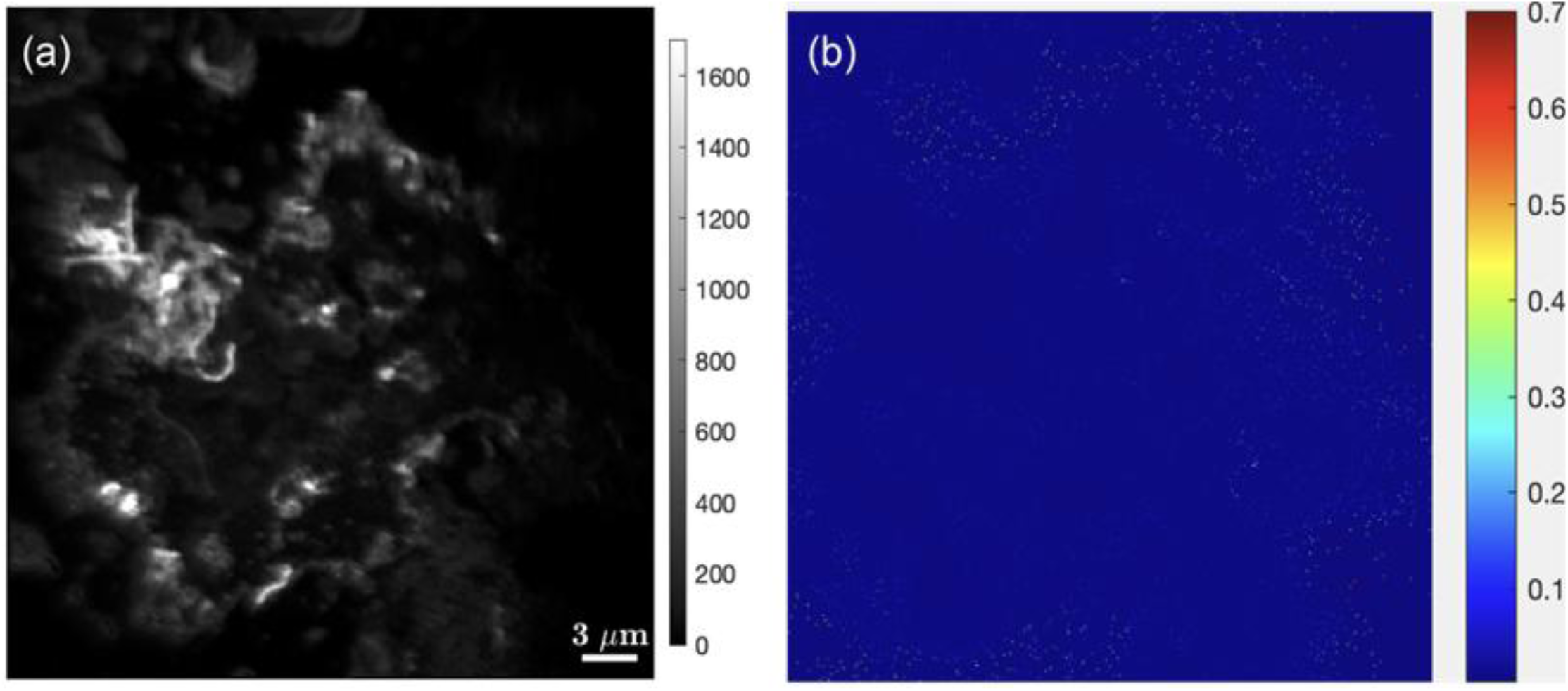
NanoSIMS image of ^15^N fractional abundance of the killed control amended with ^15^N-GlcNAc for incubated planktonic cells in electrochemical run 2 (EC2_BR3). The planktonic microbes were killed in the control culture by autoclaving at 121°C for 45 min. No enrichment of ^15^N was detected above background levels (a) nanoSIMS ion image of total biomass (^14^N^12^C^-^) showing distribution of planktonic cells in the raster image, where values in scale bar represent the pixel counts. (b) Corresponding fractional abundance ^15^N image scaled to natural abundance ^15^N (0.0036),

**Supplementary Figure 6:**
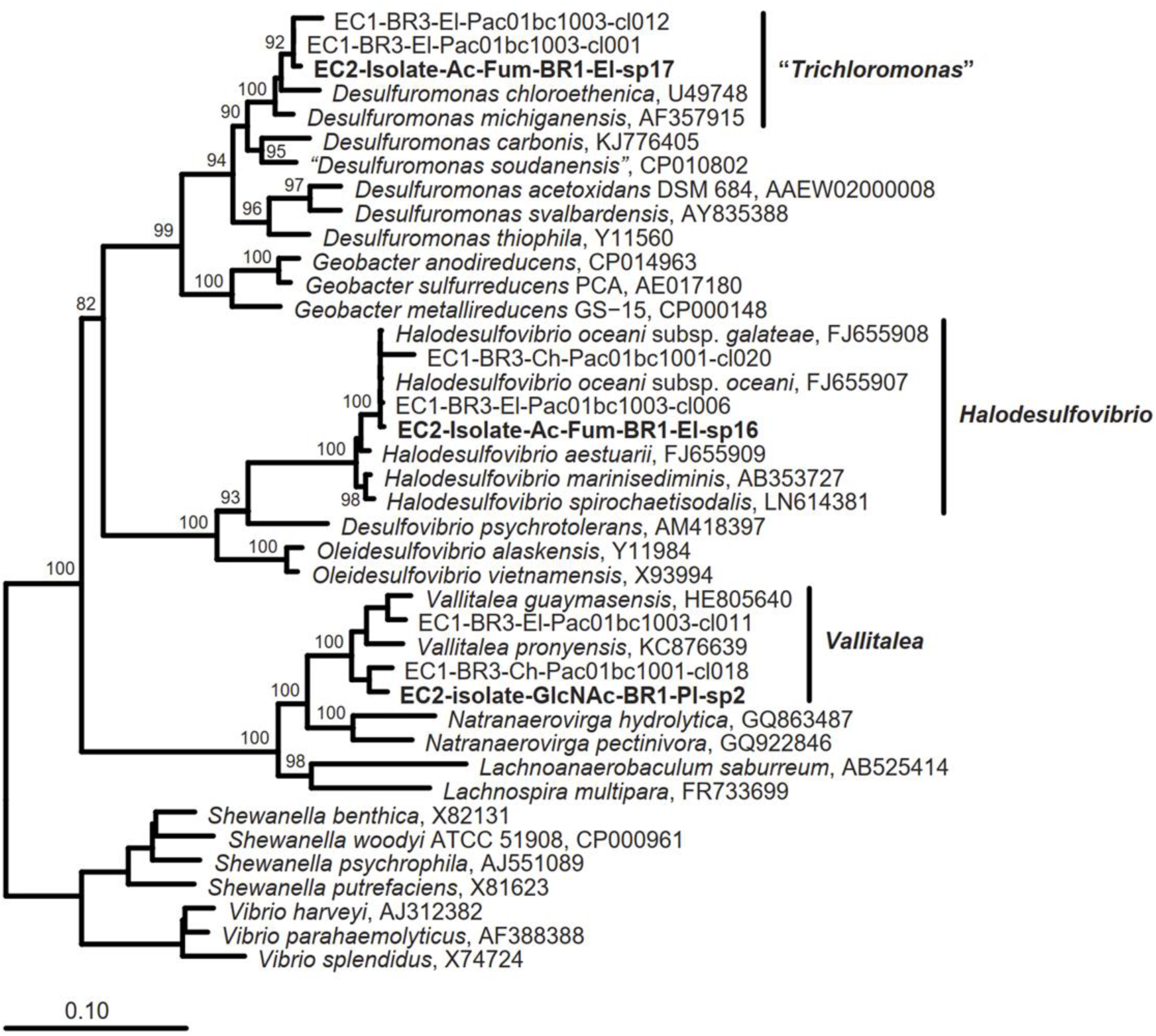
Phylogenetic tree illustrating the taxonomic affiliation of anaerobic isolates obtained from electrochemical run 2, biological replicate 1 (EC2_BR1). The isolates include *Vallitalea* sp. 2, enriched using 3 mM N-acetylglucosamine (GlcNAc), and *Trichloromonas* sp. 17 and *Halodesulfovibrio* sp. 16, enriched using 10 mM sodium acetate and 20 mM sodium fumarate. The tree was constructed using the SILVA 138.1 reference database and the ARB software (v7.1.0). The tree was generated with the Maximum Likelihood algorithm implemented in RAxML version 8, applying the GTRGAMMA substitution model. GTR rate parameters were optimized using the BFGS method. Bootstrap support values (based on 100 non-parametric replicates) are displayed for nodes with ≥85% support. Isolate sequences shown in **bold** represent shorter sequences that were incorporated into the tree using parsimony placement. Full-length 16S rRNA gene sequences obtained using PacBio from EC1_BR3, whose carbon cloth electrode (El) served as the inoculum for the EC2 reactors, are labeled with names starting with “EC1”. The tree is rooted with *Shewanella* and *Vibrio* as the designated outgroup taxa.

**Supplementary Figure 7:**
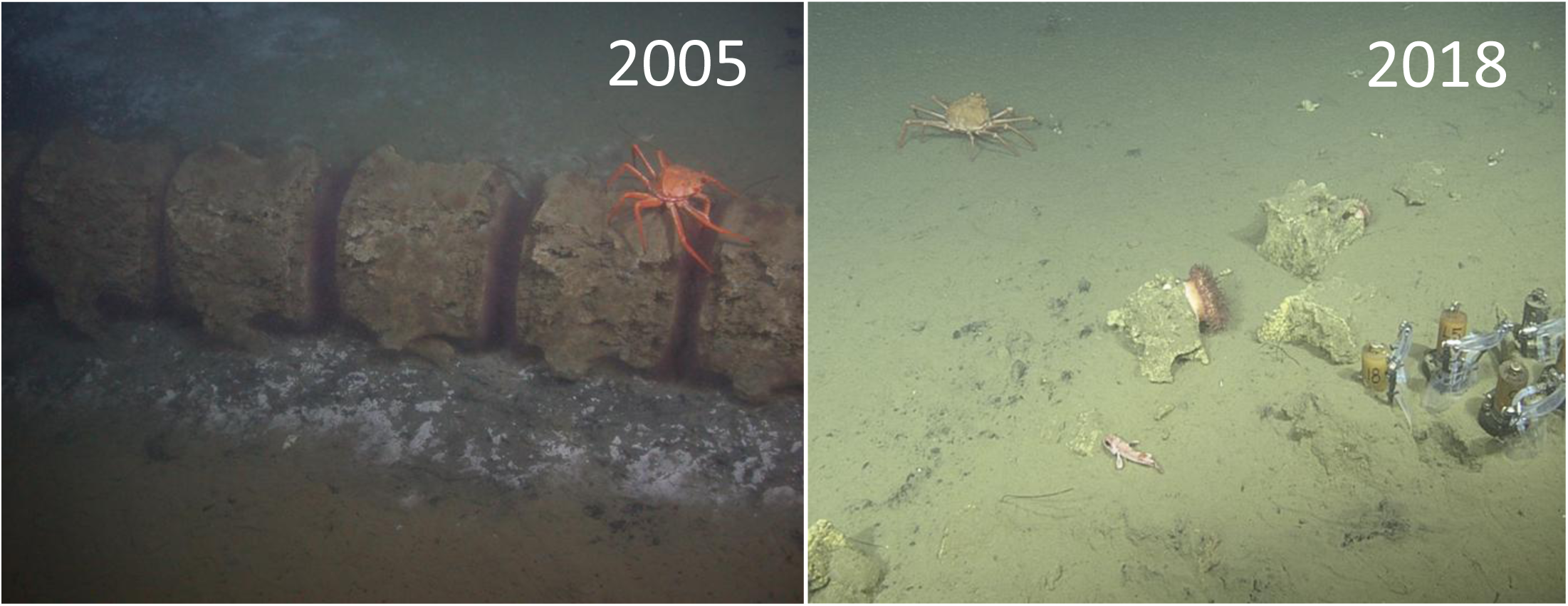
Photo taken by ROV Doc Ricketts of whalefall (WF1018) in 2005 (left) and December, 2018 (right) shows deep-sea sediment sampling at the whale fall site WF1018 in Monterey Canyon, at a depth of approximately 1018 meters. In the center of the right image, a portion of whale bone is still visible, partially embedded in the seafloor, with a sea anemone anchored nearby. This bone is part of the blue whale skeleton that continues to influence local biogeochemistry and microbial community composition nearly two decades after emplacement. Several push-core samplers are deployed using the manipulator arm of a remotely operated vehicle ROV Doc Ricketts to collect sediments for microbiological and geochemical analysis.

**Supplementary Figure 8:**
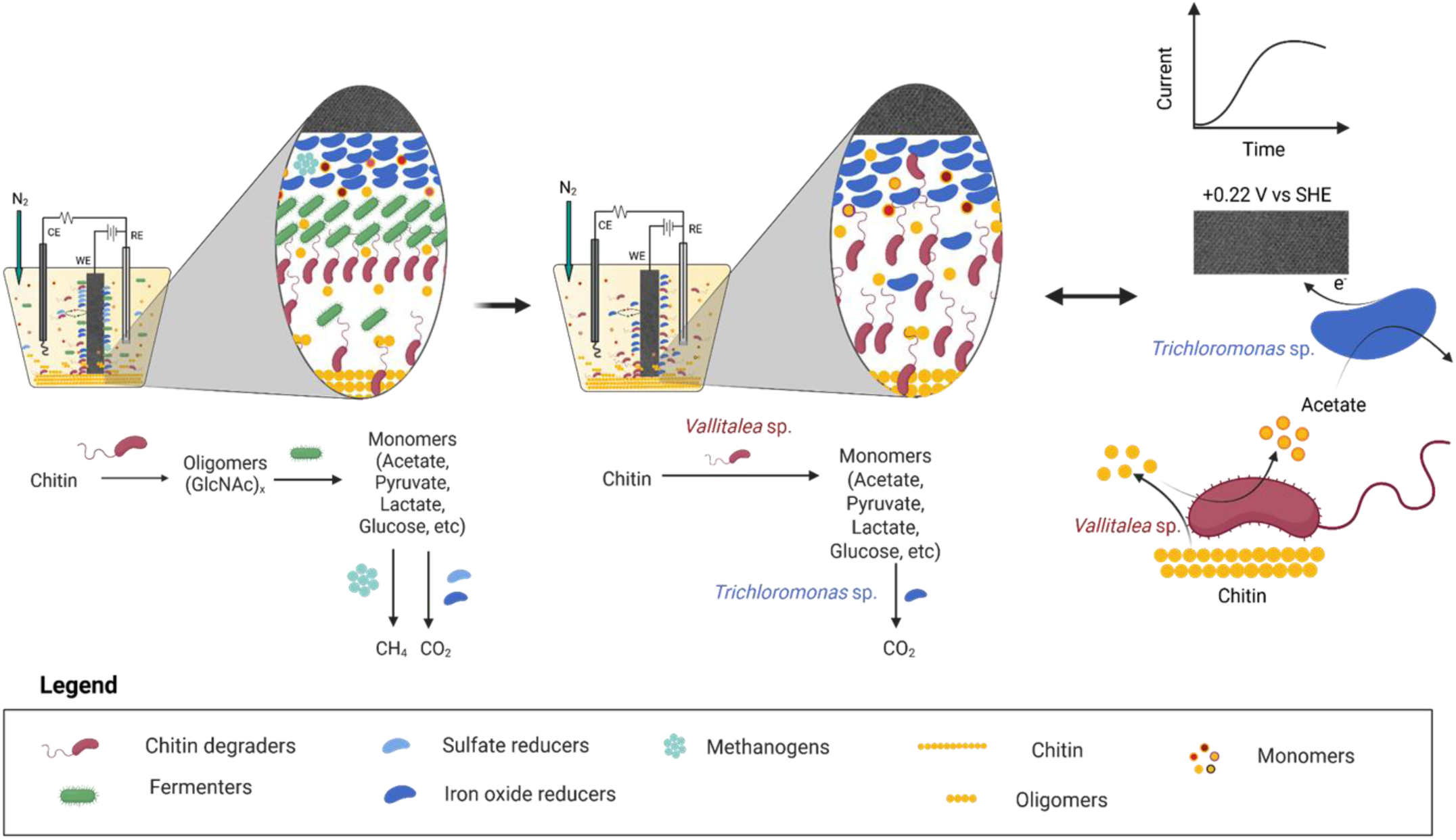
The illustration provides the spatial and temporal dynamics of microbial interactions facilitating chitin degradation and extracellular electron transfer (EET) in marine sediment-derived electrochemical incubations. Initially (left), a complex and metabolically diverse community—including chitin degraders, fermenters, sulfate reducers, iron reducers, and methanogens—mediates the degradation of insoluble chitin into oligomers (GlcNAc)x and monomers (e.g., acetate, lactate, glucose), which fuel downstream anaerobic respiration (sulfate reduction and iron oxide reduction). Upon incubation in an electrochemical reactor with a poised working electrode (WE), selective pressure under anoxic, redox-controlled conditions enriches for microbes capable of chitin degradation and extracellular electron transfer (EET). This process eventually led to the isolation of two key taxa belonging to genus *Vallitalea* and *Trichloromonas*. Reincubation of these two isolates in a minimal electrochemical setup (right) reconstituted the core syntrophic interaction, *Vallitalea* sp. (str2) catalyzes chitin degradation and fermentation, producing acetate, which is in turn oxidised by *Trichloromonas* sp. (str17) via EET to the electrode. The left-to-right transition thus reflects community succession and functional partnerships, revealing the metabolic division of labor required to couple particulate organic matter degradation with electron flow in anoxic marine deep-sea sediments.

