## SupplementaryFiles for "Deep sea anaerobic microbial community couples the degradation of insoluble chitin to extracellular electron transfer"

Affiliations

Yamini Jangir, Indian Institute of Technology, Kanpur, Uttar Pradesh, India

Sujung Lim, University of Nevada, Las Vegas, NV, USA

Fabai Wu, Easter Institute of Technology, Ningbo, China

Sammy Pontrelli, VIB-KU Leuven Center for Microbiology, Leuven, Belgium

Julia Schwartzman, Biological Sciences, University of Southern California, Los Angeles, CA, USA

Supplementary Materials

Anoxic seafloor sediment abundant in putative chitin degraders and iron reducers

A geochemical analysis of a background sediment core, located less than 2 m distance from SL12123, revealed the highest concentration of Fe (II) (0.13 mM) at a depth of 1–3 cm (see Figure 1(b)). Additionally, sediment was collected from a depth of 1–2 cm at the whale fall site WF1018 (36.771442 N, 122.082998 W) in December 2018 to probe anaerobic chitin dynamics in marine sediments.

At the community level (refer to Supplementary Figure 1 and Supplementary Table Sheet 1), analysis of nine representative (2 x SL12122, 3 x SL12123, 2 x SL-12124, 2 x SL12125) sediment samples indicated that the predominant archaeal lineage, comprising 3.1% ± 1.5%, was Bathyarchaeia within the phylum Crenarchaeota. This group, previously known as the miscellaneous Crenarchaeotal group (MCG)[1], is dominant in anoxic subsurface environments and is known for diverse carbon metabolisms, including methane cycling[2,3] In terms of bacterial lineages, the most abundant ASVs were assigned to the phyla Proteobacteria (17.7% ± 2.8%), Desulfobacterota (10.0% ± 3.5%), Planctomycetota (5.8% ± 2.4%), Acidobacteriota (5.0% ± 2.9%), Sva0485 (4.8% ± 0.8%), NB-1j (4.7% ± 1.7%), Myxococcota (3.9% ± 1.2%), Latescibacterota (3.1% ± 1.3%), Bacteroidota (2.9% ± 2.6%). Other phyla were present in lower relative abundances, including Chloroflexi (0.7% ± 0.5%), Spirochaetota (0.5% ± 1.0%), Firmicutes (0.1% ± 0.2%), and Fusobacteriota (0.1% ± 0.1%).

The dominance of gammaproteobacterial ASVs was primarily attributed to several key groups, including *Gammaproteobacteria Incertae Sedis* (5.1 ± 1.0%), BD7-8 marine group (3.5 ± 1.7%), B2M28 (3.4 ± 1.0%), *Woeseia* (2.2 ± 1.4%), uncultured *Thiohalorhabdaceae* (1.6 ± 0.7%), *Gammaproteobacteria AT s2 59* (1.4 ± 0.7%), and uncultured *Pseudomonadaceae* (0.1 ± 0.1%). Members of the *Gammaproteobacteria Incertae Sedis* have the ability to oxidize various sulfur species under anoxic conditions[4]. The term "Incertae Sedis" indicates a taxonomic group with uncertain physiology. The BD7-8 marine subgroup consists of anaerobic carbohydrate degraders, often living in symbiosis with marine sediment invertebrates[5]. B2M28, a clone first identified in seagrass-containing marine sediments, closely resembles a sulfur-oxidizing symbiont associated with the bivalve *Codakia orbicularis*[6], which thrives in marine sediments[7,8]. The genus *Woeseia*, within the order Woesiales, exhibits organoheterotrophic[9] metabolism and facultative chemolithoautotrophy[10,11], potentially enabling growth on proteinaceous substrates[12]. The family *Thiohalorhabdaceae* includes *Thiohalorhabdus*, a genus of halophilic, facultative anaerobic chemolithoautotrophs[13]. The uncultured bacterial clone *AT-s2-59*, collected from a hydrothermal vent in the mid-Atlantic Ridge, is a likely sulfur oxidizer within the *Halothiobacillus* group[14]. Additionally, ASVs were annotated to uncultured alphaproteobacterial *Rhodobacteraceae* (0.6 ± 0.4%), found broadly in marine sediments across various subgroups (excluding *Roseobacter*) but with limited physiological information[15]. Cultured representatives of this family, however, form symbiotic relationships with aquatic micro- and macroorganisms[16] and are capable of extracellular electron transfer[17]. Following Gammaproteobacteria, Desulfobacterota-associated ASVs were the most abundant. These ASVs represented taxa from the sulfate and mineral reducing Desulfobulbaceae family[18–20], putative dissimilatory iron reducer Sva1033[21–23], mixotrophs Syntrophobacterales order[24], and sulfate reducing genus Halodesulfovibrio[25,26]. Microbes from these taxa have been suggested to perform low chain fatty acid (LCFA) degradation[27,28].

The whale fall sediment also hosts ASVs annotated to the uncultivated phylum NB1-j, known for hydrocarbon degradation and prevalent in various marine environments. NB1-j may be associated with microalgae, potentially aiding in nitrogen supply[29]. Acidobacteriota (specifically Subgroup_10 and Subgroup_23) was also observed; while this lineage is predominant in soil microbiomes where it degrades polysaccharides like chitin and cellulose[30,31], it has also been detected in marine environments with potential sulfur-cycling functions on the seafloor[32]. The class *Phycisphaerae* within the order Planctomycetes, represented by MSLB9, is involved in the degradation of complex carbohydrates and is commonly found in marine sediments[33]. Within Bacteroidota (5.3 ± 1.2%), we identified an uncultured genus from the Bacteroidetes_BD2-2 group, which likely degrades proteins and amino acids in anaerobic environments[34]. This group may be associated with methanotrophic archaea and sulfate-reducing bacteria[35], particularly in methane seep sediments. However, the precise physiological roles of Bacteroidetes in sediments remain largely unresolved. The Sva0485 clade, predominantly known for sulfate and iron reduction[36], was also present, but its physiology and ecological roles remain unclear due to a lack of microbial isolates or genomes. Within the family Myxococcota (formerly Myxobacteria), we identified *Sandaracinaceae* and the *MidBa8* family. Myxobacteria, often found in marine sediments and cyanobacterial mats[37], are primarily aerobic but some can utilize alternative electron acceptors[38,39] Lastly, Latescibacterota, a clade of uncultured microbes, was associated with marine invertebrates and is capable of degrading complex polymers[40].

The microbial community structure in whale fall sediment, revealed through 16S rRNA gene analysis, highlights a rich presence of anaerobic chitin degraders and sulfate/iron reducers, making this site ideal for studying chitin degradation coupled with iron reduction. Key lineages such as *Bathyarchaeia*, *Desulfobacterota*, and various *Proteobacteria* contribute to carbon and iron cycling in anoxic conditions, supporting diverse metabolic interactions. Additionally, several uncultured groups with potentially unique metabolic roles were identified, suggesting a rich and specialized ecosystem in this deep-sea habitat.

### Laboratory incubations

The whale fall sediments were incubated in macrocosms (25 mL sediment in 60 mL serum vials) containing 0.01/mL g chitin, 0.013 g/mL PCIO, and minimal sulfate artificial seawater media. The iron incubation (PCIO_run1) included two transfers at room temperature (RT) and 10°C with three different sulfate conditions: (1) 0.2 mM sulfate, (2) 1 mM sulfate, and (3) 1 mM sulfate + 1 mM molybdate, used as a sulfur source for assimilation. Sodium molybdate was added to inhibit sulfate-reducing bacteria growth[41]. Iron reduction was assessed with a ferrozine assay[42,43]. Colorimetric ferrozine-based assay for the quantitation of iron in cultured cells at each enrichment transfer.

Microbial cultures enriched on PCIO and chitin at 10°C with 0.2 mM sulfate were selected as the inoculum for the primary electrochemical reactor. Although the initial electrochemical enrichment (labeled echem_run1, EC1) was intended to be conducted at 10°C, repeated failures of the chiller system necessitated incubation at room temperature. Three biological replicates (EC1_BR1, EC1_BR2,EC1_BR3) electrochemical reactors were established, each with working electrodes set at +0.22 V vs. SHE, for 120-day incubation. Controls included an abiotic control (EC1_AC) without inoculum and an open circuit (EC1_OC) control with only 0.2 mM sulfate provided for assimilation in all the reactors but could also be used as the electron acceptor, in this case. Planktonic phase was sampled for 16S rRNA gene analysis, external metabolites and analytes, at various intervals of time (day 78, 87, 99, and 119). EC1_BR1 showed a spike in metabolite production at day 112. Of the 22 metabolites that peaked at this time point, half were amino acids (glutamate, glutamine, threonine, valine, glycine) or intermediates of amino acid biosynthesis or degradation (Supplementary Table: EC1_BR1_amino_acids.csv). Given the relatively lower ammonium accumulation in EC1_BR1 (Figure 2d), this suggests excess nitrogen may have been expelled not only as free ammonium but also as nitrogen-containing metabolites, possibly in an effort to maintain intracellular C:N ratios. Among the biological replicates, BR3 showed the highest metabolic activity throughout the enrichment period.

To study the formation of a stable anoxic chitin-degrading community, planktonic phase (2 mL) and electrode-attached biomass (scraped from the electrode) from EC1_BR3 were used as inoculum for a secondary electrochemical incubation (echem_run2, EC2). The second electrochemical enrichment (EC2), ran from November 2019 to July 2022, a 32 month long incubation period. In the first 20 days, the medium was amended with chitin and simpler organics, such as fumarate and acetate, to facilitate the growth of putative iron oxide reducers. Over the next 300 days, sequential amendments with pyruvate, lactate, glucose, and N-acetylglucosamine (GlcNAc) resulted in similar anodic responses. By day 320, the planktonic phase was replaced with fresh chitin and fresh medium to assess the response of the electrode-attached community over three months. Current production was restored to comparable levels following the addition

of 3 mM GlcNAc. Further 16S rRNA gene sequencing, 16S rRNA FISH coupled with BONCAT and NanoSIMS, chitinase assay, external metabolites and analytes were performed to confirm anaerobic chitin degradation. Finally, two representative species from the microbial community responsible for chitin degradation and mineral reduction were isolated to establish the syntrophic interaction in the electrochemical incubation. Metadata and respective analysis for each electrochemical reactor run and samples collected is provided as supplementary files:
Supplementary_table_metadata_ch_echem_run1.xlsx,
Supplementary_table_metadata_ch_echem_run2.xlsx,
echem_run1_CA_CV_IC_exometabolites.html,
echem_run2_CA_CV_IC.html,
echem_run2_CA_chitinase_exometabolites.html,
EC1_BR1_amino_acids.csv)

### Microbial composition in laboratory incubations

The initial electrochemical enrichment (echem_run1, EC1) was performed using 5 mL of a chitin-iron enriched culture, PCIO (10 C + 1 mM sulfate, second transfer), acting as an inoculum, in triplicate reactors. The inoculum for the electrochemical incubation was dominated by members of Firmicutes, followed by Spirochaetota, Desulfobacterota, and Bacteroidota with minor representation by archaea *Methanosarcinaceae*. The samples for 16S sequencing were analysed from planktonic community after 78, 87, 99, and 119 days of chitin incubation, while the chitin-associated and electrode-associated microbial community was sampled and analysed on only 119 day of chitin incubation. Taxonomic differences among these phases were assessed using ANCOM-BC (p-adjusted method = "fdr") for the EC1_BR3. Briefly, *Gammaproteobacteria*, *Spirochaetota* (formerly grouped with Alphaproteobacteria), and *Desulfobacterota* (previously Deltaproteobacteria[18]) were detected in all three phases. Within *Gammaproteobacteria*, *Shewanella*, *Psychromonas*, and a novel *Pseudomonadaceae* genus were dominant. Notably, *Shewanella* and *Pseudomonas* are well-studied for extracellular electron transfer (EET)[44,45], while *Psychromonas* is known for biopolymer degradation under diverse conditions[46]. Within *Spirochaetota*, *Sediminispirochaeta* and *Spirochaeta_2* were enriched on chitin, while *Sphaerochaeta* was prominent in the planktonic phase. *Desulfobacterota* taxa, including *Trichloromonas* and a novel *Desulfuromonadaceae* genus, were associated with the electrode, while *Halodesulfovibrio* was more abundant on chitin but present in all phases. In the planktonic phase, *Firmicutes*, *Fusobacterota*, and *Bacteroidota* were dominant. Within *Firmicutes*, genera such as *Abyssvirga*, *Vallitalea*, and a novel *Lachnospiraceae* member were uniformly distributed. *Fusobacterota* members, including *Psychrilyobacter* (associated with marine organisms), were enriched in the planktonic phase. On the electrode, *Clostridium_sensu_stricto_7*, *Acetobacterium*, and a novel

*Clostridiaceae* member were prominent. *Cloacimonadota* and *Halobacterota* were also represented. *Cloacimonadota* was absent in the chitin-associated phase and widely distributed in the planktonic phase consistent with its metabolic versatility and predominant in anaerobic digesters[47]. Their role as acetogenic fermenters[47] has also been suggested.

The planktonic phase (2 mL) and electrode-attached biomass (scraped from the electrode) from EC1_BR3, was used as an inoculum for a secondary electrochemical incubation (EC2). The predominant microbial taxa in EC2 were consistently present across all biological replicates, inhabiting the planktonic, electrode-attached, and chitin-attached phases. The planktonic phase and chitin-attached community have higher relative abundance of *Pseudomonadaceae* and *Vallitalea* (chitin degraders and secondary consumers). On the other hand, the poised electrode was enriched with *Desulfobacterota* (mineral reducers). Certain families, in low abundances, were distributed evenly across the three phases, including *Sediminispirochaeta* and *Methanolobus*. The temporal structure of the planktonic microbial community closely followed anodic current production, which was directly influenced by deliberate modifications in the reactor's planktonic phase. Community richness gradually re-established following the removal of the initial planktonic phase and the amendment of fresh chitin. The primary contributors to richness lowering in the planktonic phase could be attributed to *Trichloromonas, Desulfuromonas, Spirochaetaceae, Abyssivirga, a novel Lachnospiraceae*, and *Shewanella*. Despite frequent sampling and replenishment of the media, the planktonic phase was repopulated quickly with *Vallitalea (Lachnospiraceae)* and a novel *Pseudomonadaceae*. Meanwhile, *Methanolobus (Methanosarcinaceae)* and *Bacteroidetes* were represented in low abundances in the planktonic phase throughout the 32 month long incubation.

A co-occurrence network for the electrode-attached biomass (EC2_BR1-3) illustrates potential relationships within this simplified microbial community, where metabolic interactions balance cooperation and competition, to sustain ecological and metabolic roles (Methods and Figure 4c). According to this analysis, the dominant archaeal lineage, *Methanolobus*, a methylotrophic methanogen, presumably consuming methanol and/or methylated compounds, is benefiting from synergistic interactions with genera *Shewanella*, *Trichloromonas*, *Desulfuromonas*, which may produce its substrates. Fermentative *Acetobacterium* and *Spirochaeta*_2, might contribute to acetate production and support Desulfuromonas. Within the network, *Shewanella*, known for its EET capability, interacts with fermenters such as *Clostridium_sensu_stricto*_7, perhaps facilitating carbon and electron flow within the system. *Lachnospiraceae*, a fermentative family, likely breaks down chitin into short-chain fatty acids, supporting *Abyssivirga* and *Desulfuromonas*. *Vallitalea* also co-occurs with many taxa, including *Trichloromonas* and *Shewanella*. *Pseudomonas*, a diverse genus known for biofilm formation, proteolytic activity, EET, and denitrification, maintains diverse interactions that may enhance community stability. In summary, within the electrochemical incubation, microbial lineages within *Desulfuromonadaceae family*, typically linked to mineral cycling, exhibited positive correlations with methanogens and fermenters, suggesting their involvement in syntrophic interactions via metabolic cross-feeding. This analysis illustrates the metabolic division of labor and revealed the dynamic interplay between a functionally partitioned microbial community, among chitin degraders, fermenters, and electron-transfer microbes, and serves as a strong example of the types of relationships occurring in whalefall sediments to sustain nutrient cycling and microbial activity.

### N-Acetyl Glucosamine (GlcNAc) metabolism

In our simplified model, the microbial community responsible for chitin degradation and GlcNAc metabolism is treated as one metabolic partner, while the EET-capable microbial community serves as the other partner. The individual half reactions with their corresponding standard Gibbs free energy change ($\Delta G_r^0$; pH 7, 25 °C , 1 bar) and actual Gibbs free energy change ($\Delta G_r$ ; pH 7.8, 22 °C, 1 bar, ionic strength of 0.7 M, and chemical composition close to the experimental composition), evaluated using the pyCHNOSZ[48] and AqEquil[49] library through WORM portal following the details provided in previous literature[50].

*GlcNAc fermentation:*

$$C_8H_{15}NO_6 + 2H_2O \rightarrow 4C_2H_3O_2^- + NH_4^+ + 3H^+ \cdots\cdots \left(\Delta G_r^0 = -39.27 kJ/mol\right)$$

$$C_8H_{15}NO_6 + 2H_2O \rightarrow 4C_2H_3O_2^- + NH_4^+ + 3H^+ \cdots\cdots (\Delta G_r = -371 kJ/mol)$$

The vast difference between $\Delta G_r^0$ and $\Delta G_r$ arises due to minimal acetate accumulation in our electrochemical reactors driving the reaction forward.

*Acetate oxidation with FeOOH (goethite) reduction:*

$$C_2H_3O_2^- + 8FeOOH + 15H^+ \rightarrow 2HCO_3^- + 12H_2O + 8Fe^{2+} \cdots\cdots \left(\Delta G_r^0 = -107.77 kJ/mol\right)$$

$$C_2H_3O_2^- + 8FeOOH + 15H^+ \rightarrow 2HCO_3^- + 8Fe^{2+} + 12H_2O \cdots\cdots (\Delta G_r = -418.67 kJ/mol)$$

Briefly, GlcNAC fermentation combined with acetate oxidation, as below, results in 8 terminal electrons. The standard gibbs free energy for the reaction at pH 7, 1 bar, and temperature 25 °C is provided below.

$$C_8H_{15}NO_6 + 6H_2O \rightarrow 3C_2H_3O_2^- + 12H^+ + 2HCO_3^- + NH_4^+ + 8e^- \cdots\cdots \left(\Delta G_r^0 = -4.6 kJ/mol\right)$$

The mathematical model estimates the concentration of acetate over time by incorporating three terms: acetate production via GlcNAc fermentation, acetate consumption for EET, and acetate assimilation for biomass synthesis. The acetate production term is expressed as $\int x\,(\frac{\partial\,[GlcNAc]}{\partial t})\,dt$ , where x represents the stoichiometric ratio of acetate molecules (1, 2, and 3) generated per GlcNAc molecule using stoichiometric balance. The acetate consumption for EET term is given by $-\int \frac{1}{CE\,.\,n_e}\left(\frac{\partial\,[e^-]}{\partial t}\right)\,dt$, where Coulombic efficiency (CE = 75%) and $n_e$=8 electrons per acetate molecule determine the efficiency of acetate oxidation to electrons. The third term, $-\int\left(\frac{\partial\,[Ac]_a}{\partial t}\right)\,dt$ , is the amount of acetate used by the microbial community for assimilation/biomass production.

Here,

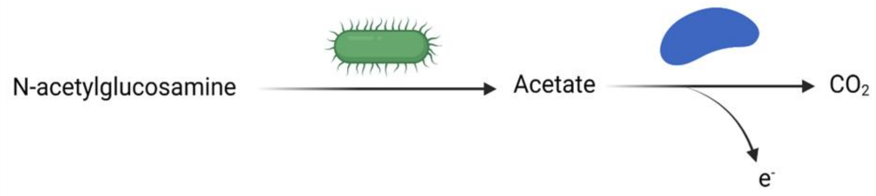


$$[Ac] = \int \frac{d\,[Ac]_+}{dt}dt - \int \frac{d\,[Ac]_-}{dt}dt$$

$$[Ac] = \int x\frac{d\,[GlcNAc]}{dt}dt - \int \frac{d\,[Ac]_{ox}}{dt}dt - \int \frac{d\,[Ac]_a}{dt}dt$$

[Ac] = acetate concentration (measured using IC),

$[Ac]_+$ = increase in acetate concentration,

$[Ac]_-$ = decrease in acetate concentration,,

x = number of acetate molecules released from GlcNAc metabolism (1 ,2, or 3) ,

[GlcNAc] = GlcNAc concentration inferred from $NH_4^-$ concentration,

CE = coulombic efficiency ,

$n_e$ = electrons per acetate molecule , and

$[e^-]$ = electron concentration.

$[Ac]_a$ = Acetate used for assimilation for biomass production

The model incorporates experimentally derived parameters, including a GlcNAc degradation rate of 0.07 mM/hr, an acetate oxidation rate of 0.0311 mM/hr via extracellular electron transfer (EET), based on a mean anodic current of 200 µA and a coulombic efficiency (CE) of 75%, and an acetate assimilation rate of 0.0032 mM/hr for biomass production. These values indicate that approximately 10% of the produced acetate is assimilated into biomass. For comparison, reported acetate uptake rates for growing *Geobacter* biofilms can[51] range from 0.014 to 0.00043 mmol $Ac^{-}$ $h^{-1}$ $cm^{-2}$. When scaled to our system (40 mL reactor volume and a minimum electrode surface area of 2 $cm^2$), these values correspond to 0.7 to 0.0215 mM/hr, which closely match the acetate oxidation rate observed in our study. As shown in Supplementary Figure 3, the time-resolved measured acetate concentrations (black dots) shows that our measured acetate concentration is a result of acetate:$NH_4^+$ stoichiometric ratio between 1:1 to 3:1. The figure illustrates an initial rise in acetate levels, followed by a decline due to acetate consumption via EET and assimilation.

Additionally (in Figure 3), we modeled the net fluxes of acetate and ammonia based on GlcNAc degradation and observed concentrations over time. Acetate flux was partitioned into EET-driven oxidation (real-time current, CE: 0.75) and assimilation (assumed at 3.2 µM/hr), while ammonia assimilation was modeled at 7 µM/hr. Based on these flux values, the acetate-to-ammonia stoichiometric ratio was calculated as approximately 1.84:1. The model was constrained by maximum acetate (15 mM) and ammonia (3 mM) produced by 3 mM GlcNAc as input.

The data and respective analysis for the above model is provided as the supplementary files and codes: G_calc_data_WORM_GB.csv,

delG_GlcNAc_fermentation.html,

sample_IC_data_current_GlcNAc.csv

flux_modelling.html
